# PI3Kδ activation, IL6 over-expression, and CD37 loss cause resistance to the targeting of CD37-positive lymphomas with the antibody-drug conjugate naratuximab emtansine

**DOI:** 10.1101/2023.11.14.566994

**Authors:** Alberto J. Arribas, Eugenio Gaudio, Sara Napoli, Charles Jean Yvon Herbaux, Chiara Tarantelli, Roberta Pittau Bordone, Luciano Cascione, Nicolas Munz, Luca Aresu, Jacopo Sgrignani, Andrea Rinaldi, Ivo Kwee, Davide Rossi, Andrea Cavalli, Emanuele Zucca, Georg Stussi, Anastasios Stathis, Callum Sloss, Matthew S. Davids, Francesco Bertoni

**Author notes:** Corresponding author: Prof. Francesco Bertoni, Institute of Oncology Research, via Francesco Chiesa 5, 6500 Bellinzona, Switzerland. Alberto J. Arribas, Eugenio Gaudio, Sara Napoli: equally contributed.

## Abstract

**Purpose:** The transmembrane protein CD37 is expressed almost exclusively in lymphoid tissues, with the highest abundance in mature B cells. CD37-directed antibody- and, more recently, cellular-based approaches have shown preclinical and promising early clinical activity. Naratuximab emtansine (Debio 1562, IMGN529) is an antibodydrug conjugate (ADC) that incorporates an anti-CD37 monoclonal antibody conjugated to the maytansinoid DM1 as payload. Naratuximab emtansine has shown activity as a single agent and in combination with the anti-CD20 monoclonal antibody rituximab in B cell lymphoma patients.

**Experimental Design:** We assessed the activity of naratuximab emtansine using *in vitro* models of lymphomas, correlated its activity with CD37 expression levels, characterized two resistance mechanisms to the ADC, and identified combination partners providing synergy.

**Results:** The anti-tumor activity of naratuximab emtansine was tested in 54 lymphoma cell lines alongside its free payload. The median IC_50_ of naratuximab emtansine was 780 pM, and the activity, primarily cytotoxic, was more potent in B than in T cell lymphoma cell lines. In the subgroup of cell lines derived from B cell lymphoma, there was some correlation between sensitivity to DM1 and sensitivity to naratuximab emtansine (r=0.28, P = 0.06). After prolonged exposure to the ADC, one diffuse large B cell lymphoma (DLBCL) cell line developed resistance to the ADC due to the biallelic loss of the *CD37* gene. After CD37 loss, we also observed upregulation of IL6 (IL-6) and other transcripts from MYD88/IL6-signaling. Recombinant IL6 led to resistance to naratuximab emtansine, while the anti-IL6 antibody tocilizumab improved the cytotoxic activity of the ADC in CD37-positive cells. In a second model, resistance was sustained by an activating mutation in the *PIK3CD* gene, associated with increased sensitivity to PI3K*δ* inhibition and a switch from functional dependence on the anti-apoptotic protein MCL1 to reliance on BCL2. The addition of idelalisib or venetoclax to naratuximab emtansine overcame resistance to the ADC in the resistant derivative while also improving the cytotoxic activity of the ADC in the parental cells.

**Conclusions:** Targeting B cell lymphoma with the CD37 targeting ADC naratuximab emtansine showed vigorous anti-tumor activity as a single agent, which was also observed in models bearing genetic lesions associated with inferior outcomes, such as MYC translocations and TP53 inactivation or resistance to R-CHOP. Resistance DLBCL models identified active combinations of naratuximab emtansine with drugs targeting IL6, PI3K*δ*, and BCL2.

Despite notable progress in recent decades, we still face challenges in achieving a cure for a substantial number of lymphoma patients (1,2). A pertinent example is diffuse large B cell lymphoma (DLBCL), the most prevalent type of lymphoma (3). More than half of DLBCL patients can achieve remission, but around 40% of them experience refractory disease or relapse following an initial positive response (3). Regrettably, the prognosis for many of these cases remains unsatisfactory despite introducing the most recent antibody-based or cellular therapies (3,4), underscoring the importance of innovating new therapeutic strategies and gaining insights into the mechanisms of therapy resistance.

CD37 is a transmembrane glycoprotein belonging to the tetraspanin family, primarily expressed on the surface of immune cells, principally in mature B cells but also, at lower levels, in T cells, macrophages/monocytes, granulocytes and dendritic cells (5) (6-8). CD37 plays a crucial role in various immune functions, including B cell activation, proliferation, and signaling, although its precise role still needs to be fully elucidated. CD37 interacts with multiple molecules, including SYK, LYN, CD19, CD22, PI3K*δ*, PI3K*γ*, and different integrins, among others (6-8). In mice, the lack of CD37 is paired with reduced T cell-dependent antibody-secreting cells and memory B cells, apparently due to the loss of CD37-mediated clustering of α_4_β_1_ integrins (VLA-4) on germinal center B cells and decreased downstream activation of PI3K/AKT signaling and cell survival (5). Reflecting the expression pattern observed in normal lymphocytes, CD37 exhibits elevated expression in all mature B-cell lymphoid neoplasms, including most lymphoma subtypes, and absence in early progenitor cells or terminally differentiated plasma cells (6,8-14). In DLBCL, CD37 expression has been reported between 40% and 90% of cases across multiple studies performed using different antibodies (10,14-16).

CD37-directed antibody- and, more recently, cellular-based approaches have shown preclinical (7,10-14,17-23) and early promising clinical activity (24-32). Among the CD37-targeting agents, naratuximab emtansine (Debio 1562, IMGN529) is an antibody-drug conjugate (ADC) that incorporates the anti-CD37 humanized IgG1 monoclonal antibody K7153A conjugated to the maytansinoid DM1, as payload, via the thioether linker, N-succinimidyl-4-(N-maleimidomethyl)cyclohexane-1-carboxylate (SMCC) (10).

Based on the initial *in vitro* and *in vivo* evidence of anti-tumor activity in lymphoma and chronic lymphocytic leukemia (CLL) (7,10), naratuximab emtansine entered the clinical evaluation as a single agent. The phase 1 study exploring naratuximab emtansine enrolled 39 patients with relapsed/refractory B cell lymphoma (27). The overall response rate (ORR) was 13% across all patients and 22% in DLBCL patients, including the only observed complete remission (CR) (27). In preliminary results of a phase 2 trial exploring the combination of naratuximab emtansine with the anti-CD20 monoclonal antibody rituximab (18), based on positive preclinical data (18), the ORR was 45% in 76 patients with DLBCL with 24 CRs (32%), 57% in 14 patients with follicular lymphoma (five CR), 50% in four MCL patients (2 CR) (31).

Here, we studied the pattern of activity of naratuximab emtansine across a large panel of cell lines derived from DLBCL and other lymphoma subtypes and characterized two resistance mechanisms to the ADC.

## Material and methods

### Cell lines

The panel of lymphoma cell lines comprised mainly models derived from DLBCL cell lines derived from DLBCL of the activated B-cell type (ABC-DLBCL, no=7) or of the germinal center B-cell type (GCB-DLBCL, no.=20), mantle cell lymphoma (no.=10), marginal zone lymphoma (no.=6), T-cell lymphomas (no.=9) and others (Supplementary Table S1).

### Compounds

Naratuximab emtansine and unconjugated toxins DM1-me were provided by Immunogen and Debiopharm International SA. Targeted agents idelalisib (PI3K*δ* inhibitor), venetoclax (BCL2 inhibitor) and S63845 (MCL1 inhibitor); monoclonal antibodies rituximab (anti-CD20) and tocilizumab (anti-IL-6R); and compounds of CHOP chemotherapy combination (cyclophosphamide, doxorubicin, vincristine and prednisolone) were purchased from Selleckchem.

### Proliferation upon single and combination treatments and Caspase 3/7 assay

Cells were seeded in 96-well plates and exposed to compounds added to create a 4-fold dilution series ranging from 200 nM to 0.19 pM and assayed by MTT following 72 hours (h) of treatment, as previously described (33). Sensitivity to single drug treatments was evaluated by the IC50 (4-parameters calculation upon log-scaled doses, R package (34) calculation. The beneficial effect of the combinations versus the single agents was considered synergism according to the Chou-Talalay combination index (35).

Apoptosis was assessed using the luminescence-base Caspase-Glo 3/7 assay kit (Promega) and defined by at least a 1.5-fold increase in signal activation concerning controls, as previously reported (33).

### CD37 and CD20 expression by FACS and by ICC

Cell lines were stained with anti-CD37 K7153A (10) (provided by Immunogen) or anti-CD20 (BioLegend #302320), and the expression was obtained by FACS, as previously described (33).

### Study of mechanism of secondary resistance

Two initially sensitive cell lines were exposed to a concentration equal to approximately 25% of the compounds’ IC50 dose, and incremental increases in dosing were performed at every passage (every 2-3 days). Parental (sensitive) were cultured parallel to resistant lines with no drug exposure. Cell identity and MDR expression were evaluated for all resistant and parental clones. The resistance was drug-specific and not MDR-mediated, and the genetic profile of all clones was confirmed (cell check9 profiling by IDEXX). For conditioned medium experiments, parental cells were cultured with 48h-conditioned or 72h-conditioned media from resistant cells, washed out in PBS, and then evaluated by the MTT proliferation assay. SU-DHL-4 parental and resistant cells were cultured for 72 hours in an exosome-depleted medium (Thermo Fisher # A2720801), washed out in PBS, and evaluated by the MTT proliferation assay. A moderated t-test (LIMMA R-package, (36)) was performed to determine statistically significant differences in drug response experiments (p<0.05). SUD-HL-4 parental and resistant lines were treated with idelalisib, venetoclax, or rituximab for 48 hours. Then, cells were permeabilized, fixed, and stained with PI (cell cycle) or double stained with Annexin V–FITC/PI (apoptosis) as previously described (37). Expression levels of CD37 were evaluated by real-time PCR.

### Whole exome sequencing and RNA-Seq

DNA and RNA sequencing were performed as previously described (38,39). Functional annotation was performed using the Gene Set Enrichment Analysis (GSEA) (40) with the Molecular Signatures Database (MSigDB)(40) and SignatureDB database (41).

### Genetic editing

Single base pair editing in the PI3KCD gene was done by homology-directed repair (HDR) using the Alt-R CRISPR-Cas9 system and HDR donor oligos (Integrated DNA Technologies, Leuven, Belgium). Cas9 single guide RNA (sgRNA) and 88 nucleotides long single-strand DNA donor template oligonucleotides were designed using the Alt-R CRISPR HDR Design Tool (https://eu.idtdna.com/pages/tools/alt-r-crispr-hdr-design-tool) and synthesized (Integrated DNA Technologies). Cells (200,000 cells per each experimental group) were nucleofected with 4D-Nucleofector (Amaxa-Lonza, Basel, Switzerland) in 20 µl of SG solution, according to manufacturer instructions and optimal protocol of nucleofection for this cell type (CM150), in the presence of RNA-protein (RNP) complex, HDR donor oligo and electroporation enhancer. In detail, the RNP complex HiFi Cas9-gRNA (Integrated DNA Technologies) was pre-assembled in a ratio of 4:4.8 µM at room temperature for 20 minutes, according to manufacturer instructions. Each electroporation was performed in the presence of 125 pmol of RNP complex, two µM of donor template oligo or control oligo, and four µM of electroporation enhancer (Integrated DNA Technologies). Cells were then cultured in the presence of 30 µM of Alt-R-HDR enhancer (IDT) for 24h. DNA was extracted, and ten ng of gDNA was tested for mutational status by allele-specific RT PCR. After one week of culture, cells underwent an MTT assay to evaluate the sensitivity to naratuximab emtansine.

### Mitochondrial priming and BH3 profiling

BH3 profiling was performed by exposing permeabilized cells to a panel of BH3-domain peptides in the presence or absence of idelalisib, as previously described (42)

### Data analyses

A moderated t-test (LIMMA R-package, (36)) was performed to determine statistically significant differences (p<0.05) in each experiment. Pearson correlation was evaluated using the R environment. The Mann-Whitney test assessed the association between IC_50_ values and genetic lesions.

## Results

### Naratuximab emtansine has strong anti-tumor activity in lymphoma cell lines

The anti-tumor activity of naratuximab emtansine was assessed across a large panel of cell lines derived from DLBCL and other lymphomas (Table 1, Supplementary Table S2). The median IC_50_ in 54 cell lines was 780 pM (95%C.I., 263 pM-11.45 nM). Activity was stronger in B cell lymphoma cell lines than in T cell lymphoma cell lines (P<0.001) (Table 2). No differences were observed among B cell lymphoma subtypes (Table 2). Among DLBCL cell lines, cell of origin (Table 2) and the presence of *BCL2* or *MYC* translocations did not affect the sensitivity to naratuximab emtansine (Supplementary Figure S1A-C). At the same time, IC50s were lower in cell lines bearing a *TP53* inactivation (P= 0.025) (Supplementary Figure S1D). The observed anti-proliferative activity of naratuximab emtansine was mainly cytotoxic. Induction of apoptosis was demonstrated in 33/54 (61%) lymphoma cell lines using a caspase 3/7 activation assay. We compared the activity of naratuximab emtansine to what we previously obtained using the *in vitro* version of R-CHOP on the same cell lines (43), and we did not observe any correlation (Supplementary Figure S2A).

**Table 1.** IC50 values (MTT upon 72hr of treatment) for naratuximab emtansine and its payload DM1. And CD37 surface levels by FACS. IC_50_ values calculated by the 4-parameters log-dose calculation, all values in pM. Median fluorescence intense levels of surface CD37 were normalized to isotype. All values from at least 2 independent experiments.

| N | Cell line | Naratuximab<br>emtansine<br>IC50 (pM) | DM1-me<br>IC50<br>(pM) | Surface<br>CD37 | N | Cell line | Naratuximab<br>emtansine<br>IC50 (pM) | DM1-me<br>IC50<br>(pM) | Surface<br>CD37 |
| --- | --- | --- | --- | --- | --- | --- | --- | --- | --- |
| 1 | SU-DHL-16 | 100 | 50 | 26 | 28 | JVM2 | 400 | 30 | 6 |
| 2 | WSUDLCL2 | 10000 | 40 | 73 | 29 | Z138 | 20000 | 40 | 2 |
| 3 | SU-DHL-8 | 40000 | 30 | 7 | 30 | GRANTA519 | 780 | 20 | 17 |
| 4 | SU-DHL-4 | 200 | 150 | 63 | 31 | JEKO1 | 30 | 12 | 4 |
| 5 | SU-DHL-10 | 50 | 25 | 29 | 32 | MAVER1 | 800 | 15 | 16 |
| 6 | DB | 500 | 100 | 115 | 33 | SP53 | 780 | 15 | 32 |
| 7 | OCI-Ly-8 | 50 | 50 | 76 | 34 | MINO | 150 | 8 | 51 |
| 8 | OCI-Ly-18 | 12000 | 40 | 54 | 35 | REC1 | 100 | 100 | 17 |
| 9 | OCI-Ly-19 | 30000 | 12 | 2 | 36 | SP49 | 12 | 12 | 49 |
| 10 | SU-DHL-5 | 25 | 15 | 21 | 37 | UPN1 | 5 | 10 | 11 |
| 11 | OCI-Ly-1 | 0.5 | 40 | 4 | 38 | KARPAS-1718 | 10 | 8 | 27 |
| 12 | DOHH2 | 30000 | 30 | 9 | 39 | SSK41 | 150 | 10 | 11 |
| 13 | KARPAS-422 | 300 | 30 | 62 | 40 | VL51 | 20000 | 40 | 8 |
| 14 | RCK8 | 40000 | 50 | 108 | 41 | ESKOL | 400 | 20 | 7 |
| 15 | VAL | 800 | 30 | 74 | 42 | HAIR-M | 250 | 40 | 5 |
| 16 | FARAGE | 40 | 10 | 67 | 43 | HC-1 | 780 | 30 | 13 |
| 17 | SU-DHL-6 | 120 | 20 | 25 | 44 | PCL12 | 12000 | 40 | 4 |
| 18 | PFEIFFER | 50000 | 40 | 118 | 45 | MEC1 | 15000 | 50 | 4 |
| 19 | OCI-Ly-7 | 50 | 50 | 101 | 46 | KARPAS-1106-P | 700 | 10 | 55 |
| 20 | TOLEDO | 15000 | 50 | 11 | 47 | FEPD | 25000 | 30 | 3 |
| 21 | SU-DHL-2 | 3000 | 40 | 51 | 48 | KARPAS299 | 40000 | 10 | 2 |
| 22 | RIVA | 150 | 30 | 9 | 49 | L82 | 20000 | 10 | 2 |
| 23 | OCI-Ly-3 | 600 | 30 | 36 | 50 | KIJK | 40000 | 5 | 2 |
| 24 | OCI-Ly-10 | 250 | 15 | 57 | 51 | SU-DHL-1 | 30000 | 8 | 1 |
| 25 | HBL1 | 50 | 50 | 6 | 52 | H9 | 15000 | 70 | 2 |
| 26 | TMD8 | 8000 | 30 | 55 | 53 | HUT78 | 13000 | 100 | 3 |
| 27 | U2932 | 15000 | 30 | 24 | 54 | MAC1 | 15000 | 5 | 2 |
|  |  |  |  |  | 55 | HH | 10000 | 200 | 4 |

**Table 2.** Anti-tumor activity of naratuximab emtansine in lymphoma cell lines. IC_50_ values obtained after 72h treatment. DLBCL, diffuse large B-cell lymphoma; ABC, activated B-cell; GCB, germinal center B-cell; MCL, mantle cell lymphoma; MZL, marginal zone lymphoma; CLL, chronic lymphocytic leukemia; PMBCL, primary mediastinal large B-cell lymphoma; SS, Sezary syndrome; ALCL, anaplastic large cell lymphoma; PTCL-NOS, peripheral T-cell lymphoma-not otherwise specified. n.d., not determined. * Upper confidence limit held at maximum of sample.

|  | No. | Median IC50<br>(pM) | 95% confidence interval<br>(pM) |
| --- | --- | --- | --- |
| B cell lymphomas | 46 | 450 | 150-800 |
| T cell lymphomas | 9 | 22,500 | 14,350-40,000 |
| ABC DLBCL | 7 | 600 | 81-12,800 |
| GCB DLBCL | 20 | 400 | 56-14651 |
| MCL | 10 | 275 | 18-794 |
| MZL | 6 | 325 | 24-18,078 |
| CLL | 2 | 13,500 | 12,000-15,000* |
| PMBCL | 1 | 700 | n.a. |
| ALCL | 5 | 30,000 | 15,000-40,000* |
| SS | 2 | 14,000 | 13,000-15,000* |
| PTCL-NOS | 2 | 12,500 | 10,000-15,000* |

In parallel, we also looked at the activity of DM1, the free payload of the ADC (Table 1). Its median IC_50_ was 30 pM (C.I.95%, 20-40 pM) with no differences between B and T cell lymphoma origin (Supplementary Table S2). Among DLBCL cell lines, no association was seen between sensitivity to DM1 with cell of origin, *TP53* status, or *BCL2* or *MYC* translocations (Supplementary Figure S3).

There was no correlation between sensitivity to DM1 and to naratuximab emtansine when we considered all the 54 B and T cell lymphoma cell lines (r=0.07, P = 0.6) (Supplementary Figure S4A). Conversely, a non-significant correlation was seen in the subgroup of cell lines derived from B cell lymphoma (r=0.28, P = 0.06) (Supplementary Figure S4B).

### *In vitro* activity of naratuximab emtansine is correlated to the expression of its target

The surface expression of CD37 was determined for all 54 cell lines by flow cytometry (Supplementary Table S1). Naratuximab emtansine IC_50_ values were inversely correlated with its target expression (Pearson correlation r=-0.32, P=0.019) (Figure 1A). The findings were corroborated using two RNA datasets we had previously obtained on the same panel of cell lines used in this study (44,45). The ADC IC_50_ values were inversely correlated with CD37 RNA levels, measured in 51 B and T cell lymphoma cell lines via a microarray-based technology (Illumina HT-12 arrays)(r=-0.5, P=0.001) (Figure 1B-C) and also when limiting the analysis to B cell lymphoma cell lines (microarray, n.=43, r=-0.33, P=0.03; RNA-Seq, n.=45, r=-0.3, P=0.01) (Figure 1D-F).

**Figure 1.**
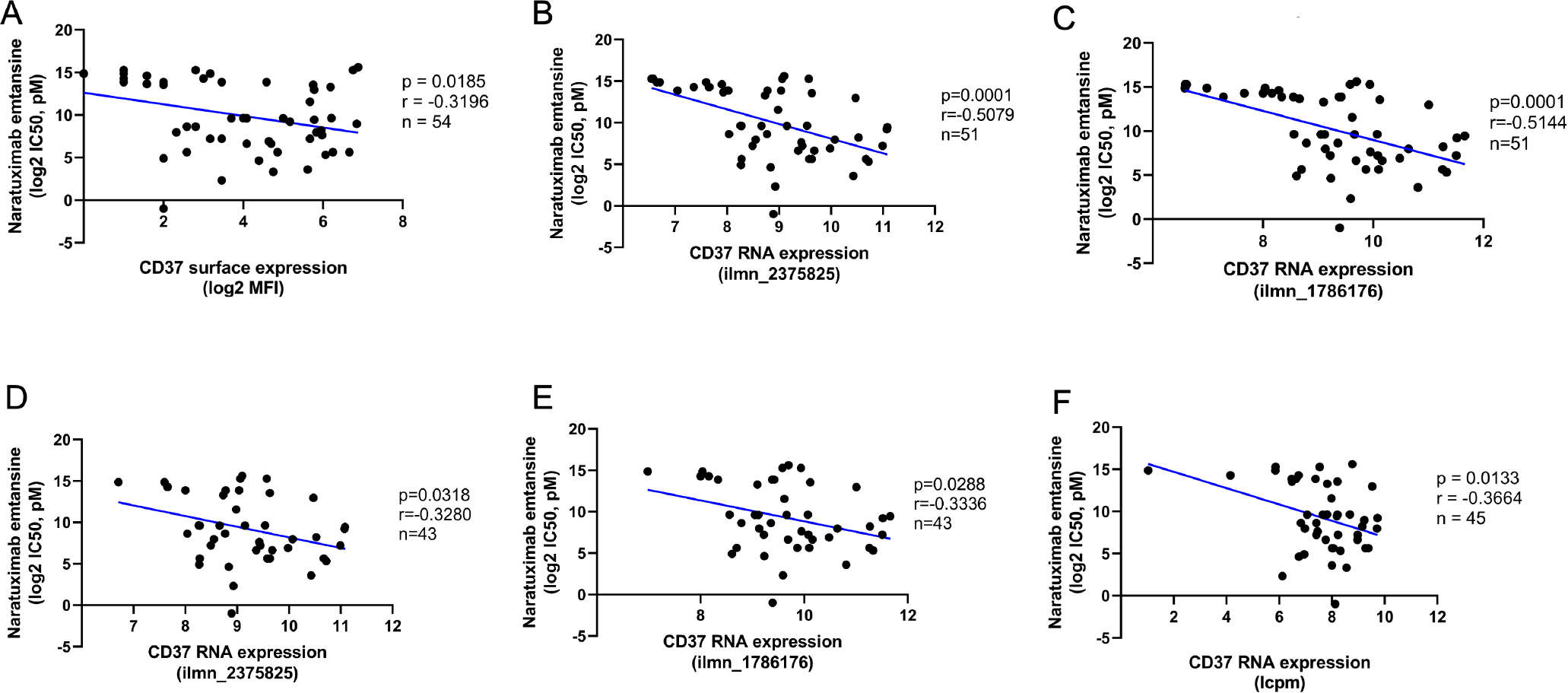
The *in vitro* cytotoxic activity of naratuximab emtansine correlated with CD37 expression. Pearson correlations between naratuximab emtansine activity, measured by IC_50_ values, with CD37 protein surface expression, measured by FACS in both B and T cell lymphomas (A, n.=54), with CD37 RNA levels, measured by the two different probes on the Illumina HT-12 arrays, in both B and T cell lymphomas (B-C, n.=51), or in B cell lymphoma only, measured with the Illumina HT-12 array (D-E, n.=43) or via total RNA-Seq in 45 B cell lymphomas (F, n.=45).

In agreement with the observed higher activity of the ADC in B than T cell lymphoma cell lines, B cell lymphoma cell lines had higher CD37 protein and RNA levels than T cell lymphomas (P<0.001) (Supplementary Figure S5A-C). No difference in CD37 expression was seen based on TP53 status in DLBCL cell lines despite the increased activity of naratuximab emtansine in this population of cells (Supplementary Figure S5E-G).

To extend the findings of the relevance of CD37 expression for the ADC’s sensitivity to the clinical setting, we first defined GCB and ABC DLBCL CD37 gene expression signatures based on the top 100 genes correlated with CD37 in DLBCL clinical specimens, and we then looked whether they were enriched in the transcripts more expressed in the DLBCL cell lines that were highly sensitive to naratuximab emtansine (IC_50_ < 800 pM) than in the resistant cell lines (IC_50_ > 10 nM). In line with what was seen for CD37 itself, Supplementary Figure S6 shows that resistant cells had lower expression of genes correlated with CD37 in DLBCL patients and that these were enriched among the transcripts higher in the sensitive cell lines.

### Development of DLBCL cell lines with resistance to naratuximab emtansine

To gain insights into potential mechanisms of resistance to naratuximab emtansine, the ADC-sensitive ABC DLBCL SU-DHL-2 and the GCB DLBCL SU-DHL-4 were exposed to increasing concentrations of the drug starting from the IC_50_ for several months until they acquired resistance to the CD37 targeting ADC. Both cell lines were also kept in culture with no drug exposure. After approximately seven months, cells kept under the drug developed resistance to naratuximab emtansine. The resistance was demonstrated to be stable by treating cells after two weeks with no drug exposure. IC_50_ values were 13-fold higher in resistant SU-DHL-2 and 6-fold higher in resistant SU-DHL-4 than in their parental cells (Figure 2A-D). Multi-drug resistance phenotype was excluded by measuring MDR1 expression via real-time PCR. Resistance was limited to the ADC and not to its payload DM1 (Figure 2B, 2D). Growth of parental cells in conditioned media taken from the resistant cells or in an exosome-depleted medium did not affect sensitivity to naratuximab emtansine (Supplementary Figure S7).

**Figure 2.**
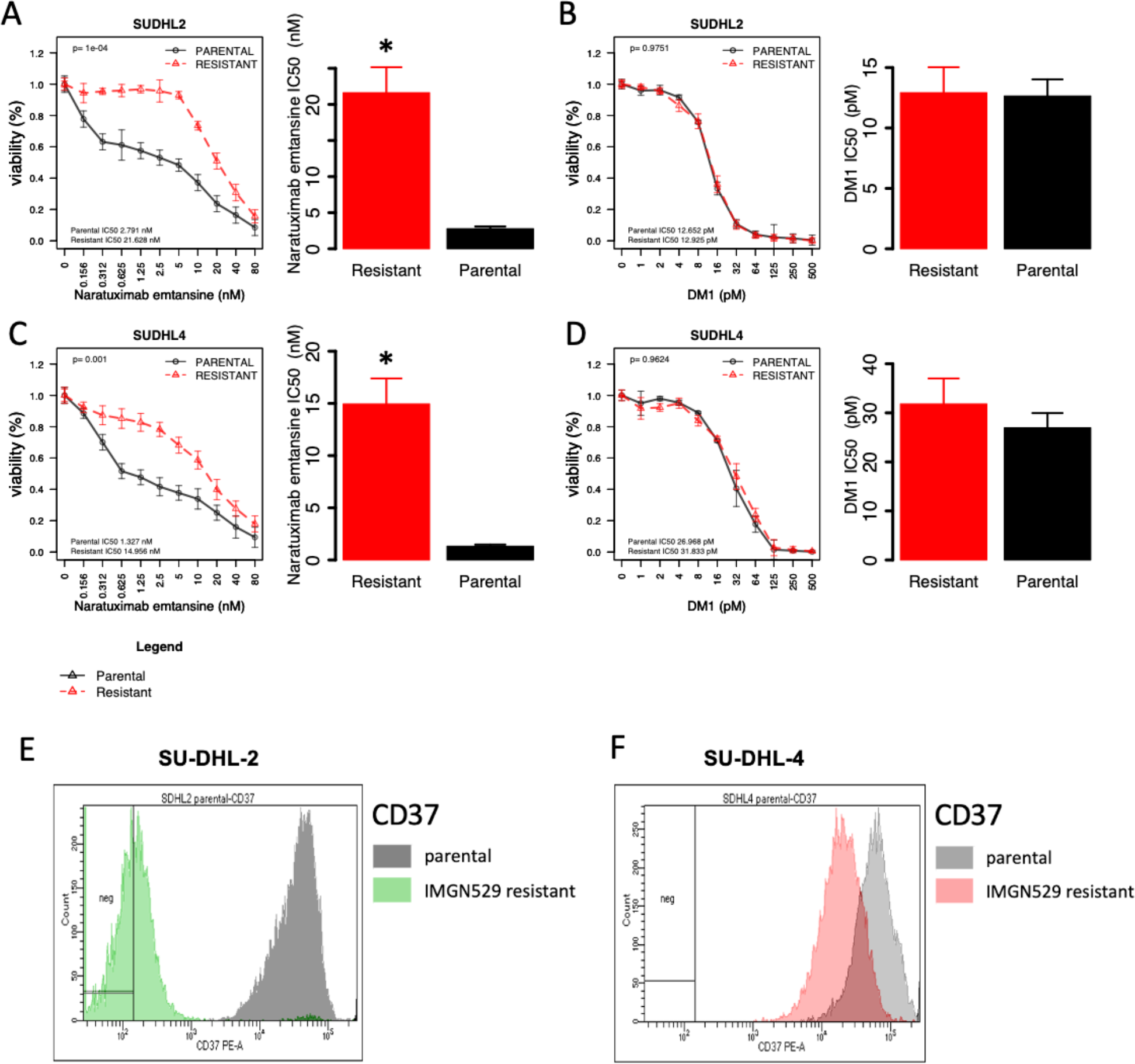
Acquired resistance to naratuximab emtansine in DLBCL cells. (A-D) MTT assay (72 h) in resistant and parental cells of SU-DHL-2 (A-B) and SU-DHL-4 (C-D) lines. Cell proliferation was evaluated in resistant and parental cells for naratuximab emtansine (Debio-1562) (left panel, A and C) and for its payload alone DM1 (right panel, B and D). Barplots correspond to IC50 values of resistant (red) and parental (black) lines. (E-F) All data correspond of at least three independent experiments. Surface protein expression of CD37 by FACS in parental and naratuximab emtansine-resistant cell lines. *, P< 0.05.

### Loss of CD37 expression due to homozygous loss as a mechanism of resistance to naratuximab emtansine

Resistant and parental cells underwent surface expression of CD37 by FACS, transcriptome analysis by RNA-Seq, and whole exome sequencing (WES) (Supplementary Table 3).

The resistant SU-DHL-2 cells developed a decrease in the surface expression of CD37 compared to their parental cells, while the reduction in the SU-DHL-4 was more limited (Figure 2). SU-DHL-4 resistant models also developed decreased expression of CD20, while SU-DHL-2 exhibited higher levels of surface CD20 (Supplementary Figure S8A-B). The SU-DHL-4 resistant cells developed reduced sensitivity also to rituximab as a single and to the combination of naratuximab emtansine plus rituximab (Supplementary Figure S8C-E).

In SU-DHL-2 cells, the downregulation of CD37 at the protein level was paired with a massive down-regulation at the RNA level (log fold change -12.68, P<0.01, FDR<0.01; (Supplementary Table S3). Whole exome sequencing and real-time PCR demonstrated a complete lack of reads mapping on the CD37 gene, compatible with the occurrence of homozygous loss (Supplementary Figure S9; Supplementary Table S3). Thus, the resistance to naratuximab emtansine in the SU-DHL-2 models could be ascribed to the loss of its target due to a genetic event.

### Activation of PI3K*δ* due to gene mutations as a mechanism of resistance to naratuximab emtansine

Transcriptome profiling in the resistant SU-DHL-4 compared to their parental cells showed enrichment of gene sets involved in PI3K signaling, lipid metabolism, and cell death (Supplementary Figure S10; Supplementary Table S3). Resistant cells exhibited de-regulation of the BCL2-family genes (Supplementary Figure S11). WES identified a series of mutations in the resistant cells, but only two variants were expressed at mRNA. One was in the *TRANK1* gene associated with a rare sleep disorder (46). The other one was represented by a heterozygous missense mutation in the *PIK3CD* gene coding for PI3K*δ* (Supplementary Table S3). The latter induced a change from asparagine (N) to threonine (T) at the PI3K*δ* amino acid 334 (N334T), and it appeared functionally relevant, being labeled as deleterious and possible damaging according to the SIFT and PolyPhen algorithms, respectively. Moreover, N334T occurred in a hotspot for mutations occurring in individuals affected by the Activated PI3K*δ* Syndrome (APDS) as well in DLBCL clinical specimens (47-51) (Supplementary Figure S12), and known to determine an increased lipid kinase activity (51).

Due to the observed BCL2 family gene deregulation and *PIK3CD* gene mutation, we exposed the SU-DHL-4 resistant cells and their parental counterpart to the BCL2 inhibitor venetoclax and the PI3K*δ* inhibitor idelalisib. The resistant cells were much more sensitive to both agents than their parental cells (Figure 3A). In both cases, we could observe an increase of apoptotic and sub-G0 cells, indicative of increased cell death induced by venetoclax and idelalisib (Figure 3B, Supplementary Figure S13 C). Each of the two agents, when given in combination with naratuximab emtansine, was able to overcome resistance to the ADC. Moreover, adding venetoclax or idelalisib to the ADC also benefited the parental cells (Figure 3C). Also, pre-treatment with idelalisib restored sensitivity to naratuximab emtansine in the SU-DHL-4 resistant cells (Supplementary Figure S13).

**Figure 3.**
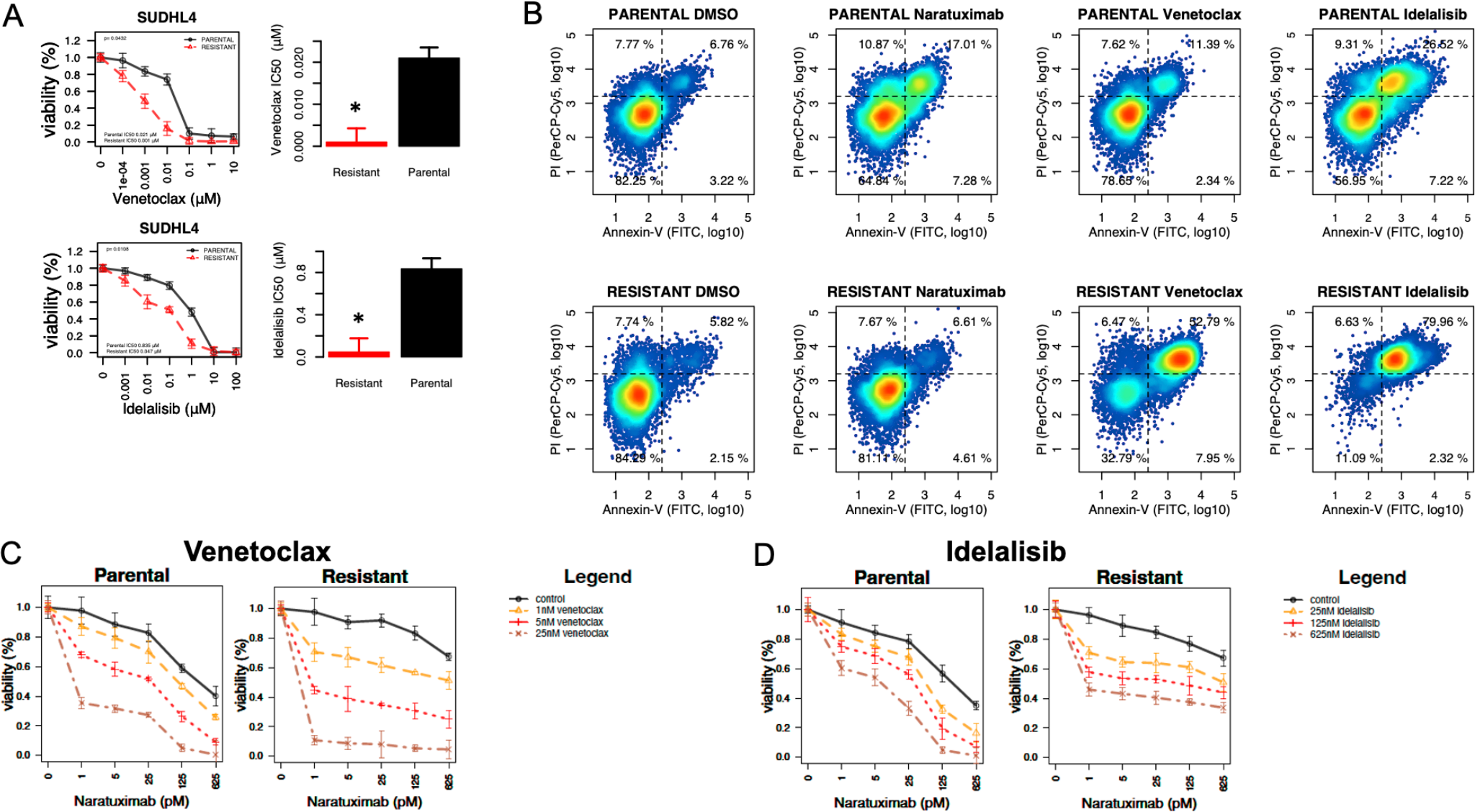
Secondary resistance to naratuximab emtansine determines increased sensitivity to idelalisib. **(A) and venetoclax (B), which can also overcome the resistance to the ADC (C). (**A) MTT assay (72 h) in resistant and parental cells of SU-DHL-4 upon increasing doses of venetoclax (upper panel) or idelalisib (bottom panel). Barplots correspond to IC_50_ values of resistant (red) and parental (black) lines. * P< 0.05. (B) Annexin-V / PI staining by FACS obtained in naratuximab emtansine resistant and parental SU-DHL-4 exposed to naratuximab emtansine (625pM), venetoclax (25 nM) or idelalisib (625nM). Representative MTT plots showing the combination of naratuximab emtansine with venetoclax (C) or idelalisib (D) in parental (left panel) and resistant (right panel) cells of SU-DHL-4. *, P< 0.05.

To better understand the functional consequences of the changes at the RNA level of members of the BCL2 family on the acquired sensitivity to venetoclax, we performed BH3 profiling on the SU-DHL-4 resistant and parental cells at baseline or in the presence of idelalisib. The cells resistant to naratuximab emtansine depended on BCL2 and BCLXL to undergo apoptosis, while parental cells relied on MCL1 and BFL1 (Figures 4A-B). Also, idelalisib increased BCL2/BCLXL dependence in the naratuximab emtansine-resistant cells but not in the parental counterpart cells (Supplementary Figure S14). These observations are consistent with the data obtained by treating the resistant and the parental cells with idelalisib. Moreover, the BH3 profiling results were further supported by exposing the SU-DHL-4 resistant and parental cells to the MCL1 inhibitor S63845 (Figure 4C). In contrast to what was observed with BCL2 inhibition, sensitivity to MCL1 inhibition was higher in parental than in the naratuximab emtansine-resistant cells (Figure 4C).

**Figure 4.**
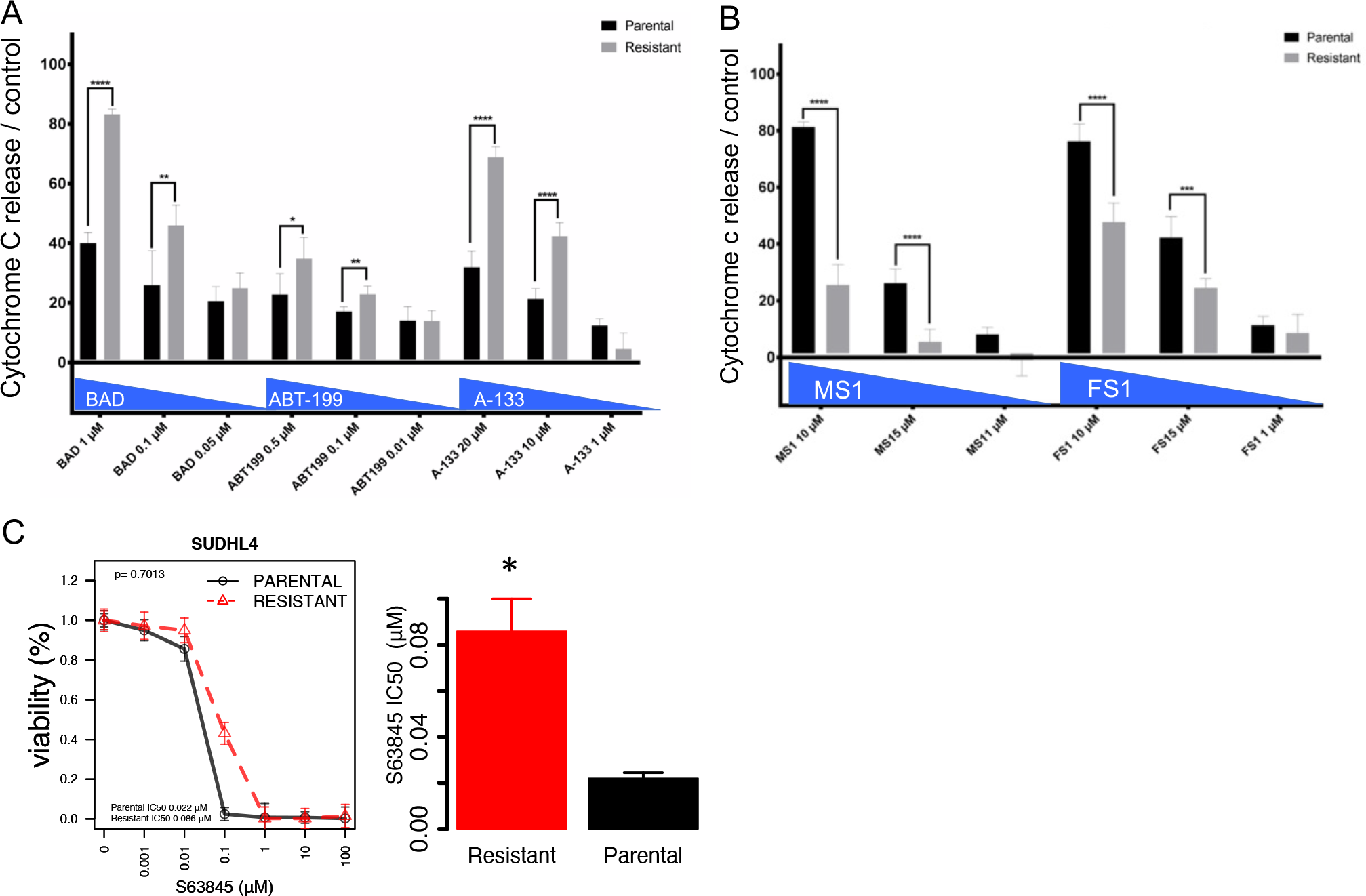
Naratuximab emtansine resistant and parental SU-DHL-4 differ based on their dependency on anti-apoptotic proteins. (A-B) BH3 profiling of naratuximab emtansine resistant (grey bars) and parental SU-DHL-4 (black bars) exploring the role of BCL2/BCLXL (BAD), BCL2 (ABT-199), BCLXL (A-133) (A), MCL1 (MS1) and BFL1 (FS1) (B). (C) MTT results obtained in naratuximab emtansine resistant and parental SU-DHL-4 exposed to increasing concentrations of the MCL1 inhibitor S63845. Barplots correspond to IC_50_ values of resistant (red) and parental (black) lines. *, P< 0.05.

Finally, we aimed to demonstrate that the *PIK3CD* N334T mutations drove resistance to naratuximab emtansine in this DLBCL model. To do this, we exploited the CRISPR Cas9 technology to create the *PIK3CD* N334T mutation (A>C, at position chr1: 9717597, GRCh38/hg38), detected in the SU-DHL-4 naratuximab emtansine resistant cells, into the genome of the SU-DHL-4 parental cells. The parental cells bearing the N334T mutation and, as control, parental cells in which we induced a silent mutation in the same locus (N334N; C>T, at position chr1: 9717598) were exposed to naratuximab emtansine. The parental cells with the *PIK3CD N334T* mutation survived the high concentration of the ADC, while the cells carrying the N334N mutations did not (Figure 5).

**Figure 5.**
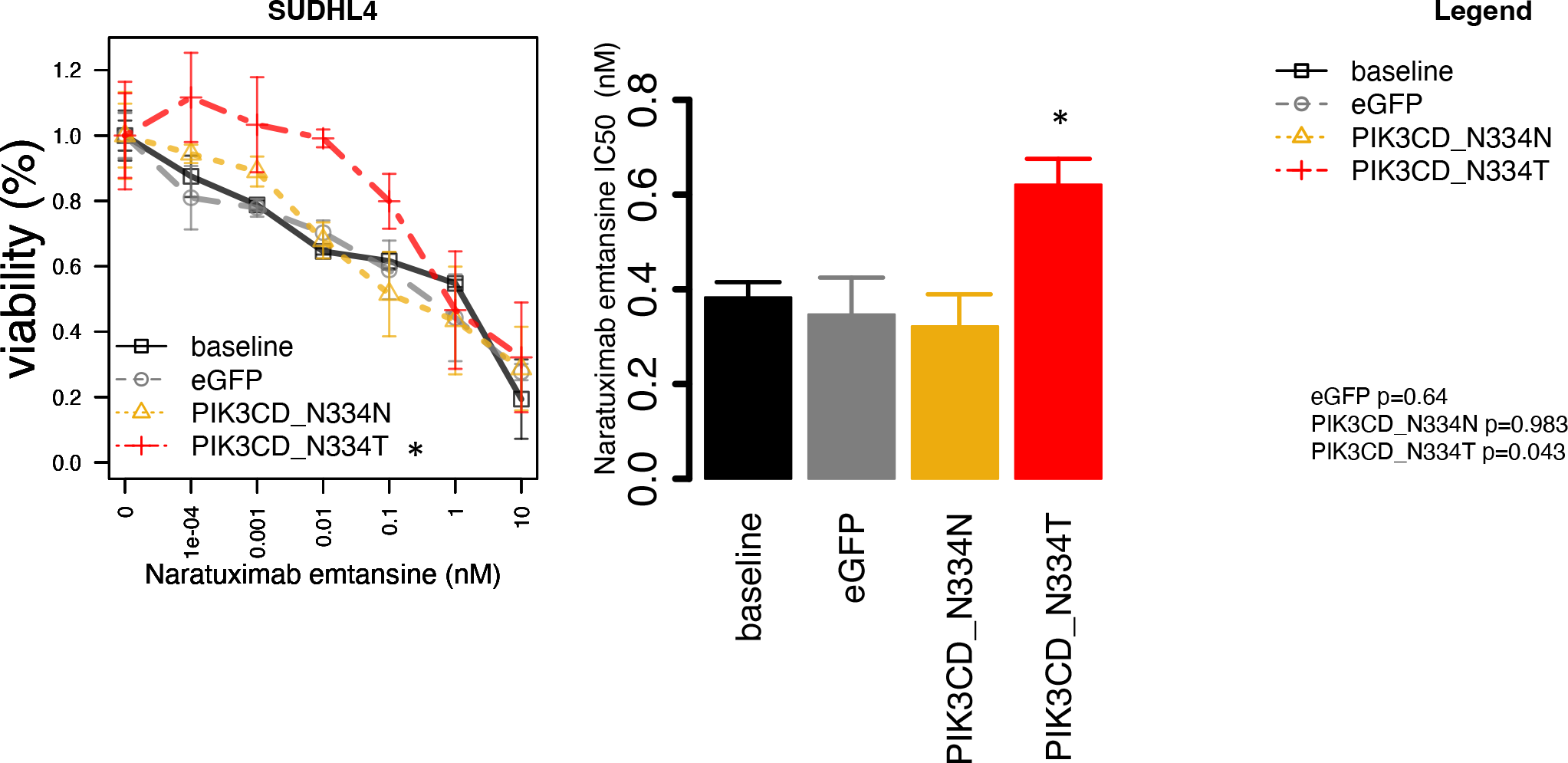
*PIK3CD N334T* mutation confers resistant to naratuximab emtansine in the SU-DHL-4 parental cells. Representative MTT results obtained in parental SU-DHL-4 cells that have undergone genome editing to induce *PIK3CD* N334T mutation (red), or, as controls, a silent mutation in the same locus (N334N) (yellow), and in parental SU-DHL-4 cells undertreated (black) or transfected with a GFP only (grey). Barplot corresponds to IC_50_ values. *, P< 0.05.

### Resistance to naratuximab emtansine does not necessarily imply sensitivity to PI3K*δ* inhibitors

We observed that the baseline transcriptome of the DLBCL cell line with resistance to naratuximab emtansine-resistant cells was enriched in the gene expression signature derived from the naratuximab emtansine-resistant SU-DHL-4 when compared to its parental counterpart (Supplementary Figure S15). Also, the transcripts higher in the resistant than in the parental SU-DHL-4 were enriched of gene expression signatures of sensitivity to PI3K inhibitors (52,53) (Supplementary Figure S16). However, when we integrated the data of sensitivity to idelalisib that we had previously obtained in the same panel of cell lines (44) and now exposed to naratuximab emtansine, we could not demonstrate a correlation between the sensitivity to the ADC and the sensitivity to the PI3K*δ* inhibitor (Supplementary Figure S17). Based on RNA-seq data (45), no mutation in the *PIK3CD* gene was present in Pfeiffer, TMD8, and MEC1, which had high naratuximab emtansine IC_50_ values and low idelalisib IC_50_ values.

### IL6 can decrease the cytotoxic activity of naratuximab emtansine

We detected an increased expression of IL6 (log fold change 2.59, P and FDR<0.01) in the CD37 negative naratuximab emtansine-resistant SU-DHL-2 (Supplementary Table S3). Transcripts more expressed in the resistant than parental cells were enriched in genes involved in cell cycle and cell proliferation and MYD88/IL6-signaling (Supplementary Table S3). Based on this observation, we assessed whether IL6 could reduce the cytotoxic activity of naratuximab emtansine. The parental and the resistant SU-DHL-2 cells were treated with the ADC and exposed to recombinant IL6, plus or minus the anti-IL6 antibody tocilizumab (Figure 6). In the parental cells, IL6 decreased the activity of naratuximab emtansine, and adding tocilizumab antagonized the effect. The latter also improved the cytotoxic effect of naratuximab emtansine in the parental cells. Due to the absence of CD37 on the cell surface, no changes were observed in the resistant cells SU-DHL-2.

**Figure 6.**
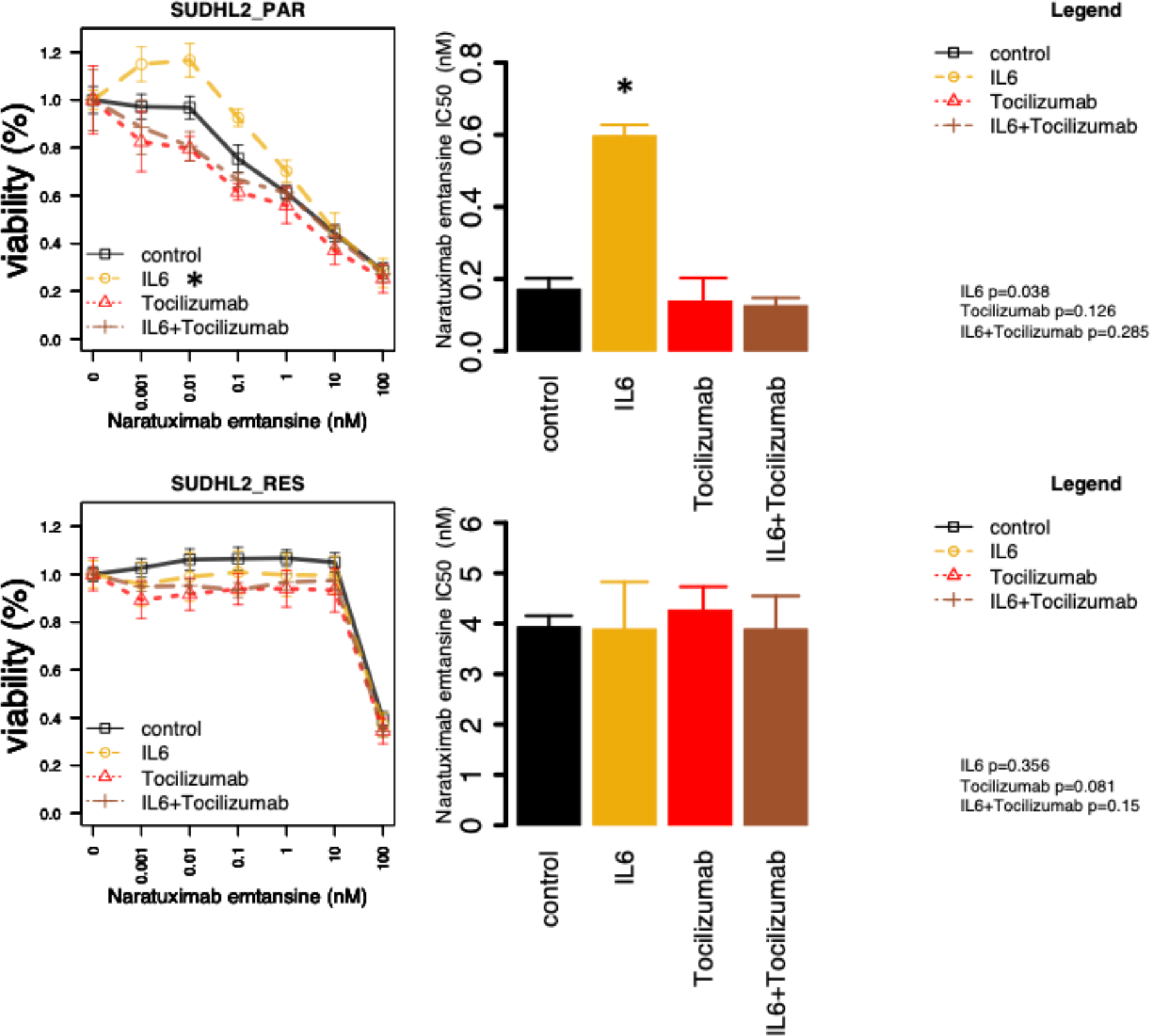
IL6 decreases the sensitivity of the DLBCL SU-DHL-2 cells to naratuximab emtansine. Parental and resistant cells were exposed (72h) to increasing concentration of naratuximab emtansine in the presence or absence of recombinant IL6 (30 ng/mL), anti-IL6 antibody tocilizumab (25µg/mL), IL6 plus tocilizumab. Cell viability was determined by MTT assay. Barplot corresponds to IC_50_ values. *, P< 0.05.

## Discussion

Here, we have characterized the anti-tumor activity of the CD37 targeting ADC naratuximab emtansine in a large panel of cell lines derived from DLBCL and other lymphomas, i) identifying subtypes that might benefit more from the treatment, ii) showing a strong correlation with the expression of its target and iii) that its pattern of activity differed from what achieved with the standard R-CHOP therapy. We also identified three potential resistance mechanisms to naratuximab emtansine, driven by the genomic loss of the gene coding for its target, acquisition of somatic mutations activating PI3K*δ*, or IL6 production.

Extending its initial description, (10), we demonstrated that naratuximab emtansine has potent *in vitro* cytotoxic activity in lymphomas, higher in B than T cell lymphomas. This aligns with the higher CD37 expression in B than T cell lymphomas, as demonstrated by our data and the literature (6,8-14). Moreover, we also observed that the *in vitro* activity of naratuximab emtansine was indeed correlated to the expression of its target, measured both at the protein level on the cell surface by FACS and at the RNA level, using different platforms.

The cytotoxic activity of naratuximab emtansine was maintained in the presence of *BCL2* or *MYC* translocations, and it was not cross-resistant with R-CHOP. The significance of the observed higher *in vitro* activity of naratuximab emtansine in TP53 inactive DLBCL cases, in the absence of a higher sensitivity to the payload and an increased expression of its target, is unclear. However, as a whole, these observations suggest that naratuximab emtansine, which has a favorable toxicity profile also in patients with severe comorbidities (54), could be explored in DLBCL patients bearing such lesions, as well as in patients refractory to or relapsed after R-CHOP who are not CAR T cell candidates.

To understand potential mechanisms of resistance that could occur in patients treated with naratuximab emtansine or with other anti-CD37 directed therapies, we have kept two DLBCL cell lines under drug pressure for many months, leading to two different models of resistance to naratuximab emtansine, that nonetheless retain sensitivity to the payload DM1. The two models also showed resistance to combining the ADC with rituximab, which has promising early clinical data (31).

One model, derived from the ABC DLBCL SU-DHL-2, was sustained by the biallelic loss of the *CD37* gene. This observation indicates that lymphoma cells can survive without CD37 expression and agrees with the report of CD37 loss of expression in up to 60% of DLBCL patients (9) and with a possible tumor suppressor role for CD37 itself (55). In mice, the lack of CD37 drives lymphomagenesis, inducing the constitutive activation of the IL6 signaling pathway (55). Interestingly, after CD37 loss, we also observed an upregulation of IL6 and other transcripts from MYD88/IL6 signaling. Moreover, we demonstrated that even IL6 alone could give resistance to naratuximab emtansine and that adding the anti-IL6 antibody tocilizumab improves the cytotoxic activity of the ADC in CD37-positive cells. Our data further identify IL6 as a resistance mechanism to multiple treatments (38,56,57). Also, they suggest the benefit of regimens including drugs that interfere with IL6 signaling and/or production (58-61) for CD37 negative lymphomas with high IL6 production (55) and for CD37 positive in combination with CD7 targeting agents.

The second model of resistance, derived from the GCB DLBCL SU-DHL-4, was sustained by the acquisition of a somatic mutation in the *PIK3CD* gene coding for PI3K*δ* and a transcriptome profile, indeed, characterized by an enrichment of gene sets related to PI3K signaling, and dysregulation of the BCL2-family genes. The *PIK3CD* N334T mutation was annotated as functionally relevant, it occurred in a hotspot for mutations occurring in individuals affected by the Activated PI3K*δ* Syndrome (APDS) (50,51) as well as in clinical specimens both from human (47,48) and canine DLBCL (49), and it was functionally proven to give resistance to naratuximab emtansine when introduced in the otherwise naratuximab emtansine-sensitive parental SU-DHL-4 cells. Moreover, the resistant derivative had also become more sensitive to PI3K*δ* inhibition and BCL2 inhibition than its parental counterpart. The latter could be explained by the switch from MCL1 dependence in the parental cells to BCL2 dependence in the resistant derivative, as demonstrated functionally by BH3 profiling. Notably, adding idelalisib or venetoclax to naratuximab emtansine overcame resistance to the ADC in the resistant derivative. It improved the cytotoxic activity of the ADC in the parental cells. Similar results have been obtained by combining another CD37 targeting agent, the CD37 antibody BI 836826, with idelalisib (62,63). All these results suggest that the combination of naratuximab emtansine with BCL2 inhibitors or with a new generation of PI3K*δ* inhibitors might be worthy of clinical evaluation in B cell lymphoma, particularly in DLBCL.

The clinical relevance of somatic mutations leading to PI3K*δ* activation in leading to resistance to naratuximab emtansine and possibly to other CD37 targeting antibodies-based therapies will require the analysis of primary samples from clinical trials. We showed that the cell lines most resistant to naratuximab emtansine in our initial screen did not carry *PIK3CD* mutations and that there was no correlation between sensitivity to the ADC or to idelalisib across 34 B cell lymphoma cell lines exposed to both compounds.

In conclusion, targeting B cell lymphoma with the CD37 targeting ADC naratuximab emtansine showed pronounced anti-tumor activity as a single agent, which is evident also in models bearing genetic lesions associated with inferior outcomes, such as MYC translocations and TP53 inactivation, or resistance to R-CHOP. Our DLBCL resistance models identified active combinations of naratuximab emtansine with drugs targeting IL6, PI3K*δ*, and BCL2.

## Authors contributions

AJA developed resistant lines, performed experiments, analyzed and interpreted data, performed data mining, prepared figures, and co-wrote the manuscript; EG performed experiments, analyzed and interpreted data; SN performed genetic editing and interpreted data; CJYH performed BH3 profiling; CT performed experiments; RPB, GS performed flow-cytometry analyses; LC performed data mining; NM performed data mining and prepared figures; LA analyzed and interpreted data; JS performed structural modeling of the PI3K*δ* complex; AR performed genomics experiments; IK performed data mining; AC performed structural modeling of the PI3K*δ* complex; EZ, AS, DR: provided advice and edited the manuscript CS and MD codesigned research and edited the manuscript; FB designed the study, interpreted data, and co-wrote the manuscript; all authors approved the final manuscript.

## Conflict of interests

**Alberto J. Arribas:** travel grant from Astra Zeneca, consultant for PentixaPharm. **Eugenio Gaudio**: Floratek employee. **Charles Jean Yvon Herbaux:** honoraria from AbbVie, Janssen, Roche, Gilead, Takeda, Daiichi; personal fees from AbbVie, Janssen, Roche, Gilead, Takeda; research funding from AbbVie, Takeda. **Chiara Tarantelli**: travel grant from iOnctura. **Luciano Cascione**: travel grant from HTG. Ivo Kwee: **Anastasios Stathis**: institutional funding for clinical trials from Debiopharm, Innomedica, Abbvie, ADC Therapeutics, Amgen, Astra Zeneca, Bayer, Cellestia, Incyte, Loxo Oncology, Merck MSD, Novartis, Pfizer, Philogen, Roche: institutional funding for consultant/expert testimony/advisory board: Debiopharm, Janssen, AstraZeneca, Incyte, Eli Lilly, Novartis, Roche, Loxo Oncology; travel grant: Incyte; Astra Zeneca. **Emanuele Zucca**: advisory boards of BeiGene, BMS, Curis, Eli/Lilly, Incyte, Janssen, Merck, Miltenyi Biomedicine and Roche; received research support from AstraZeneca, Beigene, BMS/Celgene, Incyte, Janssen, and Roche, received travel grant from BeiGene, Janssen, Gilead, and Roche. **Davide Rossi**: honoraria from AstraZeneca, AbbVie, BeiGene, BMS/Celgene, Janssen; research funding from AstraZeneca, AbbVie, BeiGene, Janssen. **Georg Stussi**: travel grants from Novartis, Celgene, Roche; consultancy fee from Novartis; scientific advisory board fees from Bayer, Celgene, Janssen, Novartis; speaker fees from Gilead. **Callum M. Sloss**: ImmunoGen employee. **M.S. Davids** has received institutional research funding from AbbVie, AstraZeneca, Ascentage Pharma, Genentech, MEI Pharma, Novartis, Surface Oncology, TG Therapeutics and personal consulting income from AbbVie, Adaptive Biosciences, Aptitude Health, Ascentage Pharma, AstraZeneca, BeiGene, BMS, Celgene, Curio Science, Eli Lilly, Genentech, Genmab, Janssen, Merck, Nuvalent, Secura Bio, TG Therapeutics, and Takeda. **Francesco Bertoni**: institutional research funds from ADC Therapeutics, Bayer AG, BeiGene, Floratek Pharma, Helsinn, HTG Molecular Diagnostics, Ideogen AG, Idorsia Pharmaceuticals Ltd., Immagene, ImmunoGen, Menarini Ricerche, Nordic Nanovector ASA, Oncternal Therapeutics, Spexis AG; consultancy fee from BIMINI Biotech, Helsinn, Menarini; advisory board fees to institution from Novartis; expert statements provided to HTG Molecular Diagnostics; travel grants from Amgen, Astra Zeneca, Beigene, iOnctura. The other Authors have nothing to disclose.

The current affiliation for Eugenio Gaudio is at Floratek Pharma, Aubonne, Switzerland, for Charles Jean Yvon Herbaux at Centre Hospitalier Universitaire de Montpellier, Montpellier, France, and for Ivo Kwee at BigOmics Analytics SA, Lugano, Switzerland.

---

**Supplementary Table S1.** Lymphoma cell lines histology and doubling time. DLBCL, diffuse large B-cell lymphoma; GCB, germinal center B-cell; ABC, activated B-cell; MCL, mantle cell lymphoma; MZL, marginal zone lymphoma; CLL, chronic lymphocytic leukemia; PMBCL, primary mediastinal large B-cell lymphoma; SS, Sezary syndrome; ALCL, anaplastic large cell lymphoma; PTCL, peripheral T-cell lymphoma not otherwise specified.

| N | Cell line | Histology | Cell of origin | Doubling time (hours) |
| --- | --- | --- | --- | --- |
| 1 | SU-DHL-16 | DLBCL | GCB | 40 |
| 2 | WSUDLCL2 | DLBCL | GCB | 62 |
| 3 | SU-DHL-8 | DLBCL | GCB | 48 |
| 4 | SU-DHL-4 | DLBCL | GCB | 39 |
| 5 | SU-DHL-10 | DLBCL | GCB | 40 |
| 6 | DB | DLBCL | GCB | 50 |
| 7 | OCI-Ly-8 | DLBCL | GCB | 24 |
| 8 | OCI-Ly-8 | DLBCL | GCB | 36 |
| 9 | OCI-Ly-19 | DLBCL | GCB | 31 |
| 10 | SU-DHL-5 | DLBCL | GCB | 35 |
| 11 | OCI-Ly-1 | DLBCL | GCB | 28 |
| 12 | DOHH2 | DLBCL | GCB | 44 |
| 13 | KARPAS422 | DLBCL | GCB | 65 |
| 14 | RCK8 | DLBCL | GCB | 36 |
| 15 | VAL | DLBCL | GCB | 60 |
| 16 | FARAGE | DLBCL | GCB | 48 |
| 17 | SU-DHL-6 | DLBCL | GCB | 37 |
| 18 | PFEIFFER | DLBCL | GCB | 24-30 |
| 19 | OCILY7 | DLBCL | GCB | 19 |
| 20 | TOLEDO | DLBCL | GCB | 43 |
| 21 | SU-DHL-2 | DLBCL | ABC | 24 |
| 22 | RIVA | DLBCL | ABC | 48-72 |
| 23 | OCI-Ly-3 | DLBCL | ABC | 30 |
| 24 | OCI-Ly-10 | DLBCL | ABC | 95 |
| 25 | HBL1 | DLBCL | ABC | 48 |
| 26 | TMD8 | DLBCL | ABC | 24-30 |
| 27 | U2932 | DLBCL | ABC | 49-50 |

| N | Cell line | Histology | Doubling time (hours) |
| --- | --- | --- | --- |
| 28 | JVM2 | MCL | 48-70 |
| 29 | Z138 | MCL | 24 |
| 30 | GRANTA519 | MCL | 48 |
| 31 | JEKO1 | MCL | 26-50 |
| 32 | MAVER1 | MCL | 24-48 |
| 33 | SP53 | MCL | 57 |
| 34 | MINO | MCL | 50-72 |
| 35 | REC1 | MCL | 38 |
| 36 | SP49 | MCL | 60 |
| 37 | UPN1 | MCL | NA |
| 38 | KARPAS1718 | MZL | 35-50 |
| 39 | SSK41 | MZL | 30-45 |
| 40 | VL51 | MZL | 70-80 |
| 41 | ESKOL | MZL | NA |
| 42 | HAIR-M | MZL | 48 |
| 43 | HC-1 | MZL | 45 |
| 44 | PCL12 | CLL | 40-50 |
| 45 | MEC1 | CLL | 40 |
| 46 | KARPAS1106-P | PMBCL | 60-100 |
| 47 | FEPD | ALCL | 56 |
| 48 | KARPAS299 | ALCL | 30 |
| 49 | L82 | ALCL | 40 |
| 50 | KIJK | ALCL | 50-72 |
| 51 | SU-DHL-1 | ALCL | 40-50 |
| 52 | H9 | SS | 27-38 |
| 53 | HUT78 | SS | 26-65 |
| 54 | MAC1 | PTCL-NOS | 30-50 |
| 55 | HH | PTCL-NOS | 50-70 |

**Supplementary Table S1.**
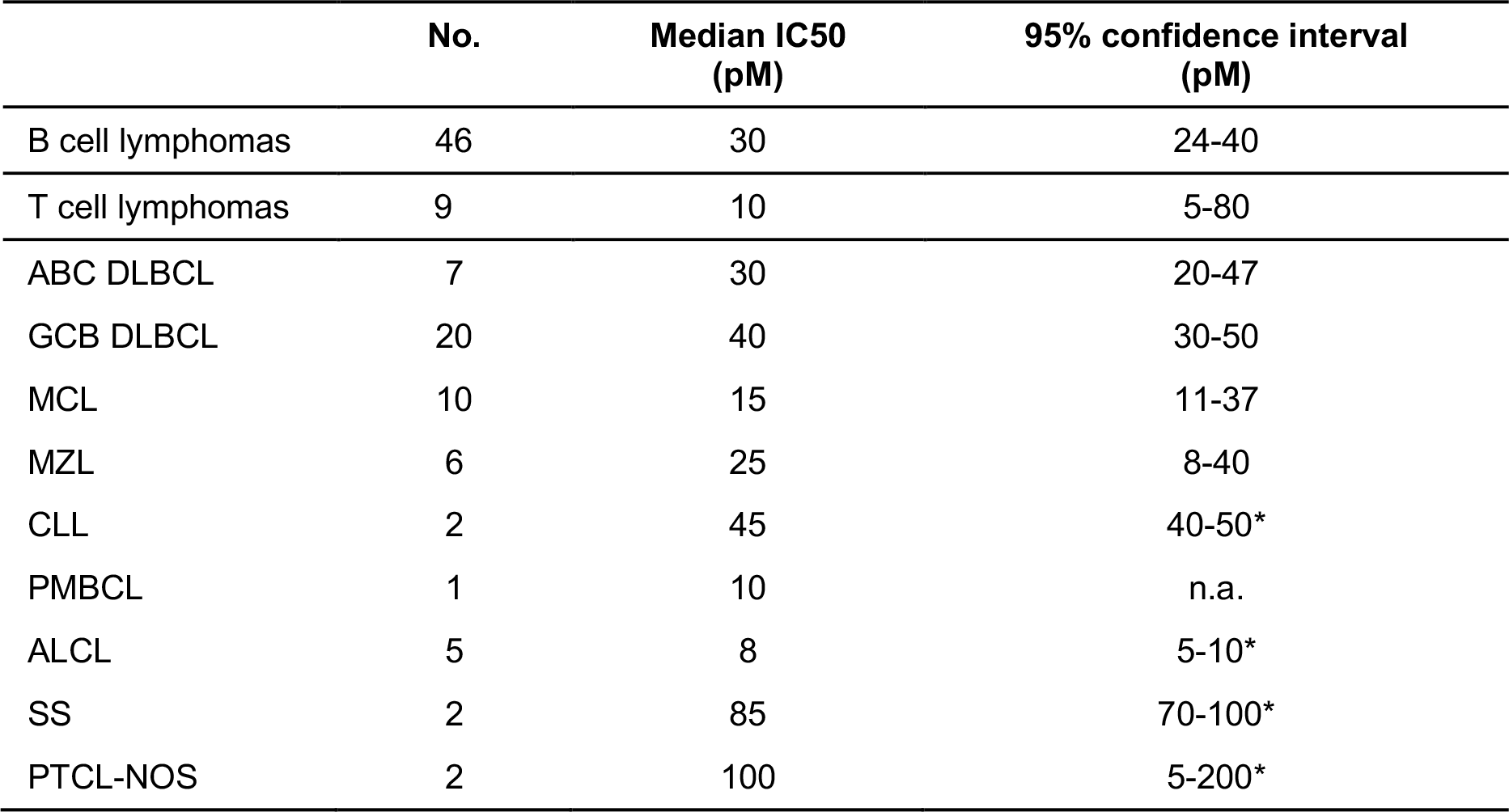
Anti-tumor activity of the DM1 payload in lymphoma cell lines. IC50s obtained after 72h treatment. DLBCL, diffuse large B-cell lymphoma; ABC, activated B-cell; GCB, germinal center B-cell; MCL, mantle cell lymphoma; MZL, marginal zone lymphoma; CLL, chronic lymphocytic leukemia; HL, Hodgkin lymphoma; PMBCL, primary mediastinal large B-cell lymphoma; SS, Sezary syndrome; ALCL, anaplastic large cell lymphoma; PTCL-NOS, peripheral T-cell lymphoma-not otherwise specified. n.d., not determined. * Upper confidence limit held at maximum of sample.

|  | No. | Median IC50<br>(pM) | 95% confidence interval<br>(pM) |
| --- | --- | --- | --- |
| B cell lymphomas | 46 | 30 | 24-40 |
| T cell lymphomas | 9 | 10 | 5-80 |
| ABC DLBCL | 7 | 30 | 20-47 |
| GCB DLBCL | 20 | 40 | 30-50 |
| MCL | 10 | 15 | 11-37 |
| MZL | 6 | 25 | 8-40 |
| CLL | 2 | 45 | 40-50* |
| PMBCL | 1 | 10 | n.a. |
| ALCL | 5 | 8 | 5-10* |
| SS | 2 | 85 | 70-100* |
| PTCL-NOS | 2 | 100 | 5-200* |

**Supplementary Table S3.** (Excel file). Transcriptome profiles by RNA-seq of SUDHL2 or SUDHL4 resistant compared to its corresponding parental lines (moderated t-test). Single nucleotide variant calling (SNV) and copy number variant calling (CNV) of of SUDHL2 or SUDHL4 resistant cells. Variants already present in the corresponding parental lines were filtered out.

