## supplementary figures and tables for "PI3Kδ activation, IL6 over-expression, and CD37 loss cause resistance to the targeting of CD37-positive lymphomas with the antibody-drug conjugate naratuximab emtansine"

**Supplementary Figure S1. Distribution of $IC_{50}$ values of naratuximab emtansine among DLBCL cell lines based on the presence of absence of *BCL2* and *MYC* chromosomal translocations, as single or concomitant events (double hit) and of *TP53* status.** A) DLBCL cell lines with (n.=15) and without (n.=11) *BCL2* translocation. B) DLBCL cell lines with (n.=10) and without (n.=16) *MYC* translocation. C) DLBCL cell lines with (n.=7) and without (n.=19) concomitant *BCL2* and *MYC* translocation. D) DLBCL cell lines with (n.=15) and without (n.=8) *TP53* inactivation. *, P< 0.05 as determined by Mann-Whitney test.

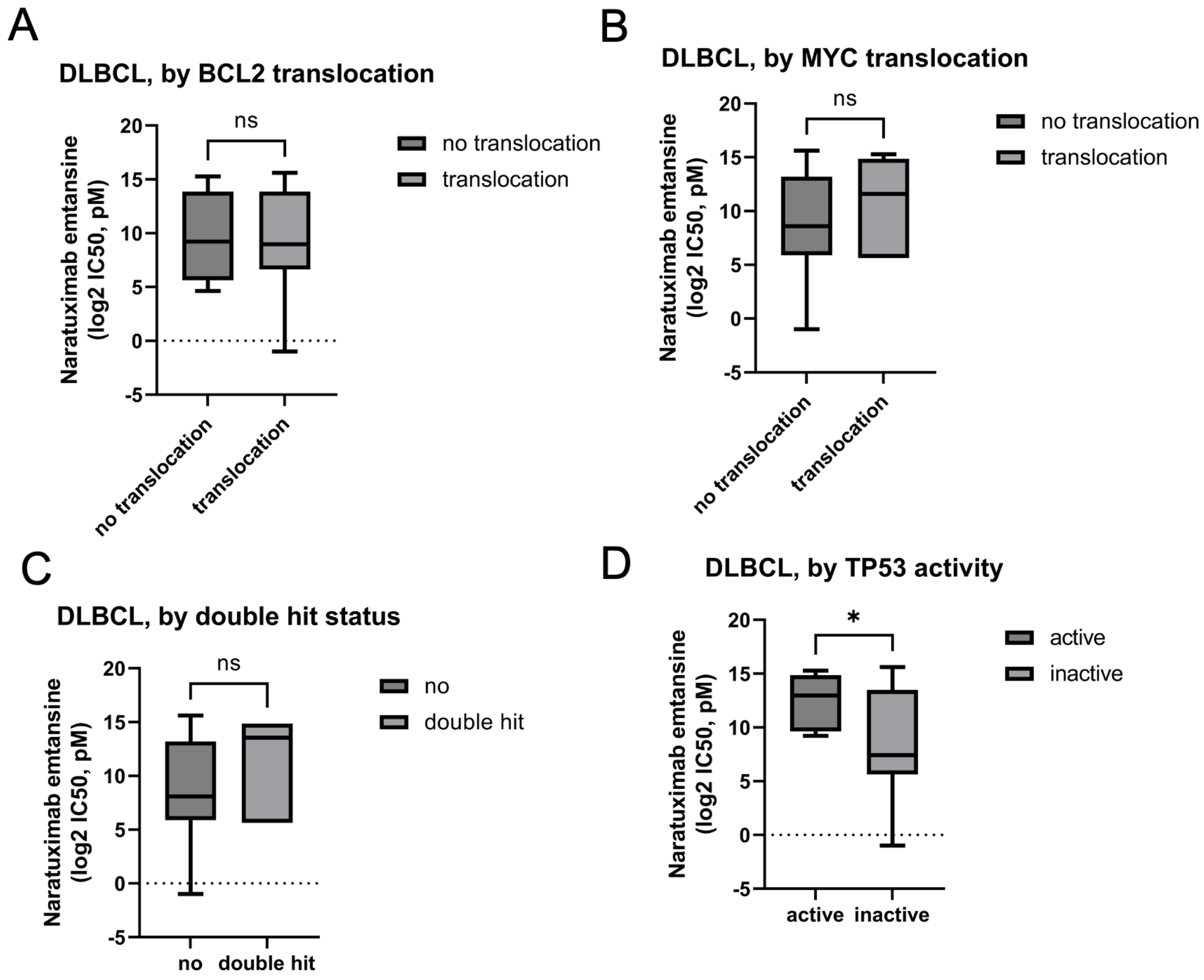

**Supplementary Figure S2. Correlation between the activity of naratuximab emtansine or its payload DM1 and R-CHOP in DLBCL cell lines.** Pearson correlations between R-CHOP and naratuximab emtansine (A, n.=37) or DM1 (B, n.=27).

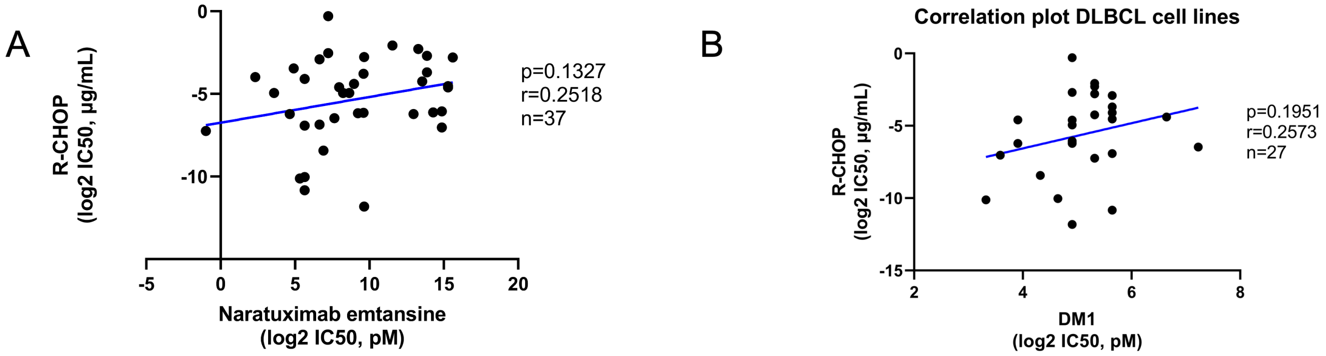

**Supplementary Figure S3. Distribution of $IC_{50}$ values of DM1 among DLBCL cell lines based on the presence of absence of *BCL2* and *MYC* chromosomal translocations, as single or concomitant events (double hit) and of *TP53* status.** A) DLBCL cell lines with (n.=15) and without (n.=11) *BCL2* translocation. B) DLBCL cell lines with (n.=10) and without (n.=16) *MYC* translocation. C) DLBCL cell lines with (n.=7) and without (n.=19) concomitant *BCL2* and *MYC* translocation. D) DLBCL cell lines with (n.=15) and without (n.=8) *TP53* inactivation. *, P< 0.05 as determined by Mann-Whitney test.

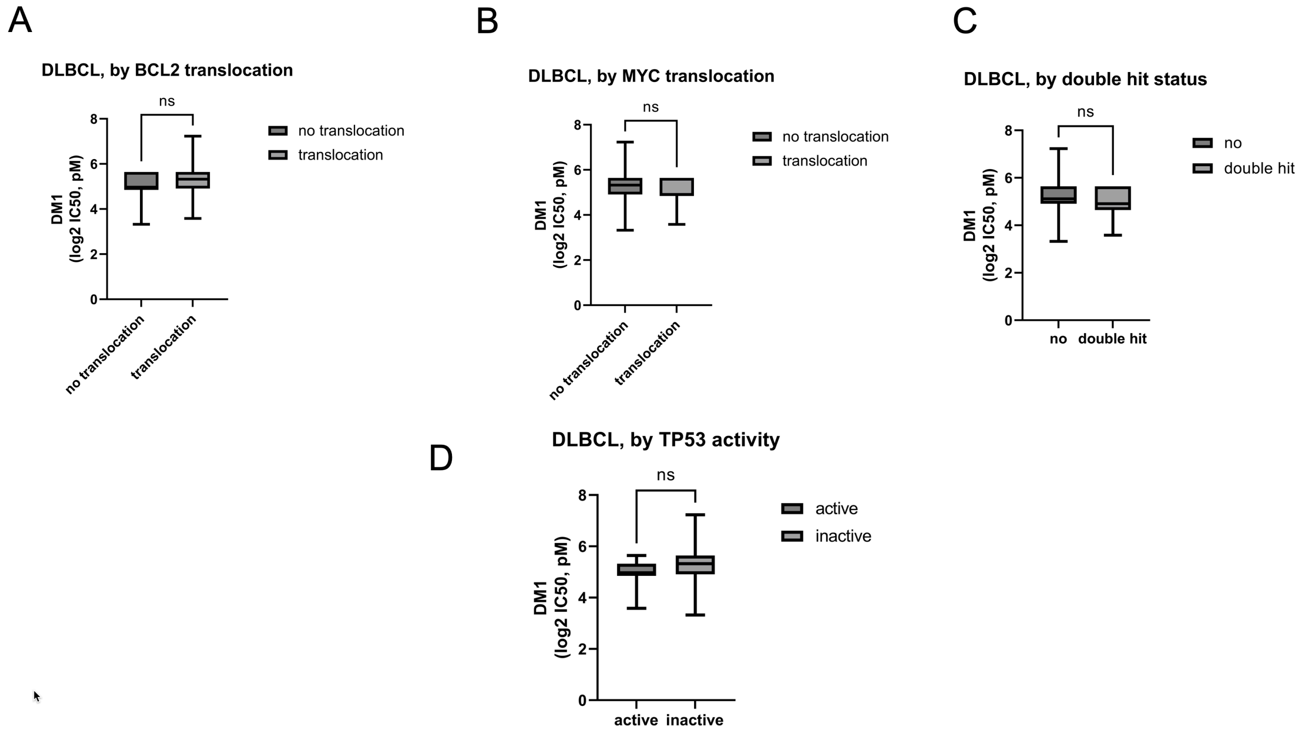

**Supplementary Figure S4. Correlation between the activity of naratuximab emtansine and its payload DM1.** Pearson correlations between log2 $IC_{50}$ obtained with naratuximab emtansine or with DM1 in 54 lymphoma cell lines (A, n.=54) and in the subset of 46 B cell lymphoma cell lines (B, n.=46).

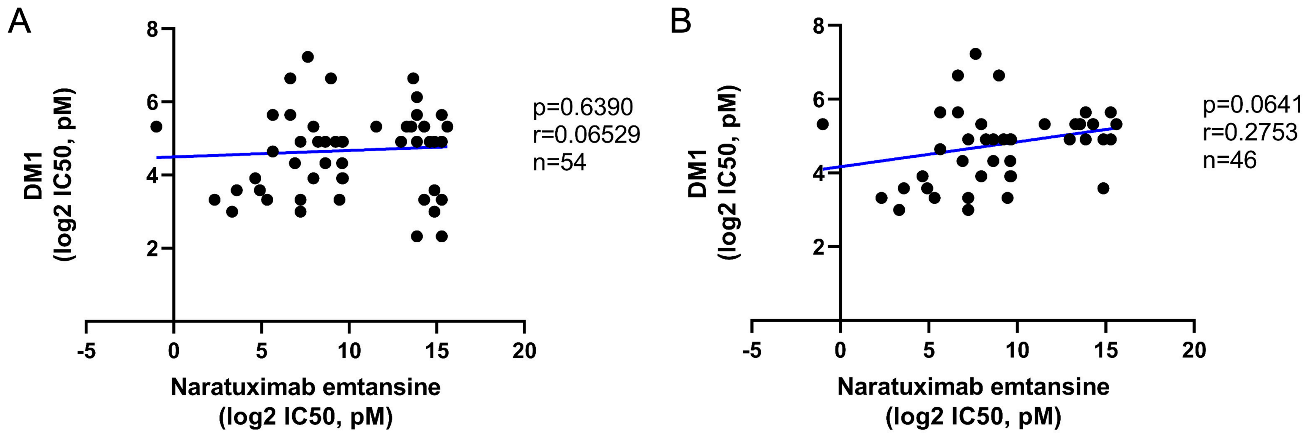

**Supplementary Figure S5. Distribution of CD37 expression based on B and T origin (A-C) and on *TP53* status (D-G).** A) CD37 surface expression between B and T cell lymphoma cell lines. B-C) CD37 RNA expression values measured via microarray between B (n.=46) and T (n.=8) cell lymphoma cell lines. D) CD37 protein surface expression, measured by FACS in DLBCL cell lines with and without *TP53* inactivation. E-F) CD37 RNA levels, measured by the two different probes on the Illumina HT-12 arrays, in DLBCL cell lines with and without *TP53* inactivation. G) CD37 RNA levels, measured via total RNA-Seq, in DLBCL cell lines with and without *TP53* inactivation. ****, P<0.0001 as determined by Mann-Whitney test.

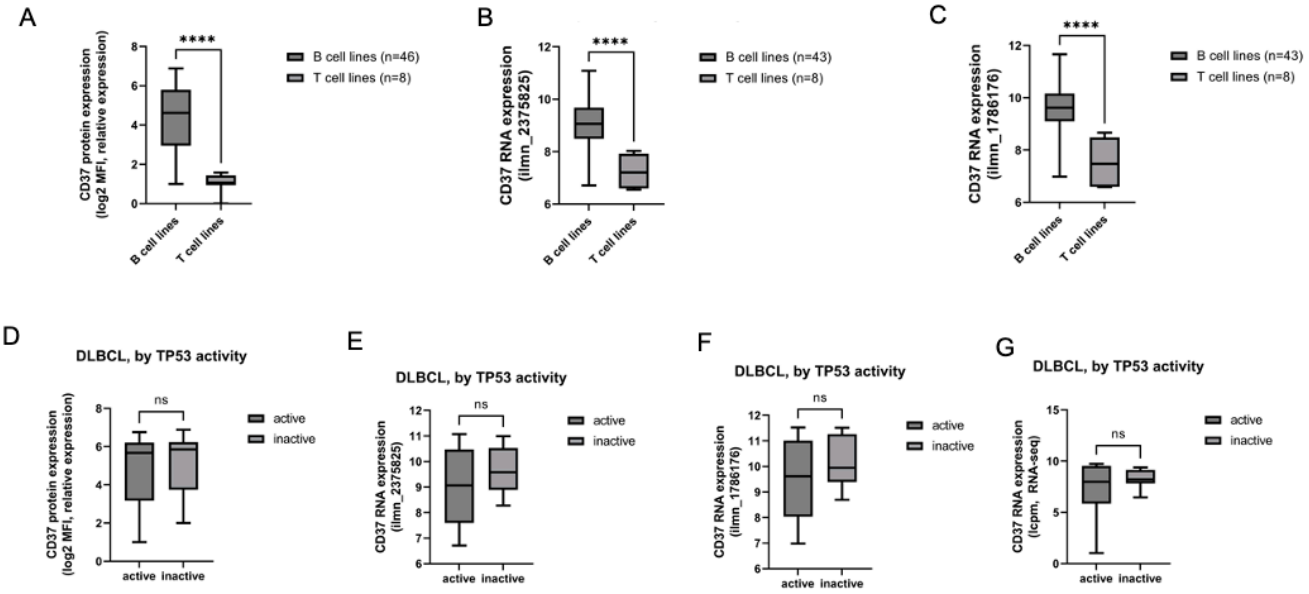


**Supplementary Figure S6. Sensitivity to naratuximab emtansine is associated to a CD37 gene expression signature.** GSEA plots for CD37 gene expression signatures derived from GCB (left plot) and from ABC (right plot) DLBCL and then analyzed for their enrichment in baseline gene-expression profiles of DLBCL cell lines that were highly sensitive to naratuximab emtansine (IC50 < 800 pM) compared to the resistant cell lines (IC50 > 10 nM). Green line, enrichment score; bars in the middle portion of the plots show where the members of the gene set appear in the ranked list of genes; Positive or negative ranking metric indicates, respectively, correlation or inverse correlation with the profile; NES, normalized enrichment score.

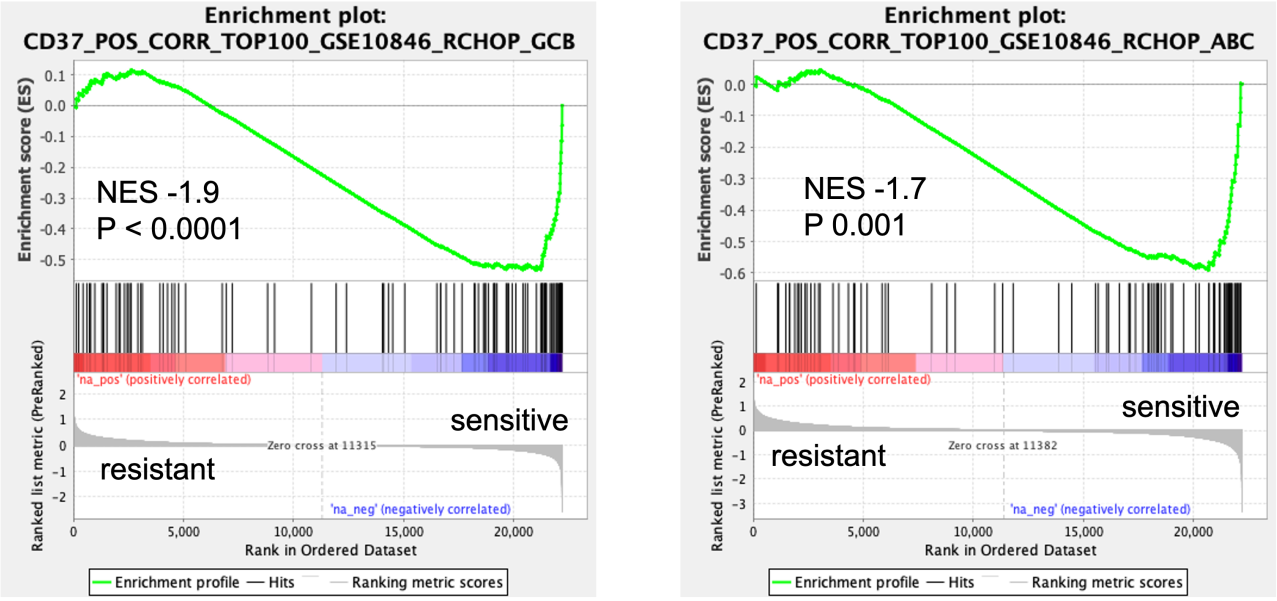

**Supplementary Figure S7. Culture of parental cells with conditioned medium from resistant cells or exosome-depleted media do not affect sensitivity to SU-DHL-4**. (A) SU-DHL-4 parental and resistant cells were cultured with resistant-conditioned medium for 48hr or 72hr (A) or cultured with exosome-depleted medium for 72hr (B). Then cells were washed out in PBS and tested for sensitivity to naratuximab emtansine. Cell viability was determined by MTT assay upon 72 hr of drug exposure. Barplots correspond to $IC_{50}$ values of resistant (red) and parental (black) lines. * P< 0.05. *, P< 0.05.

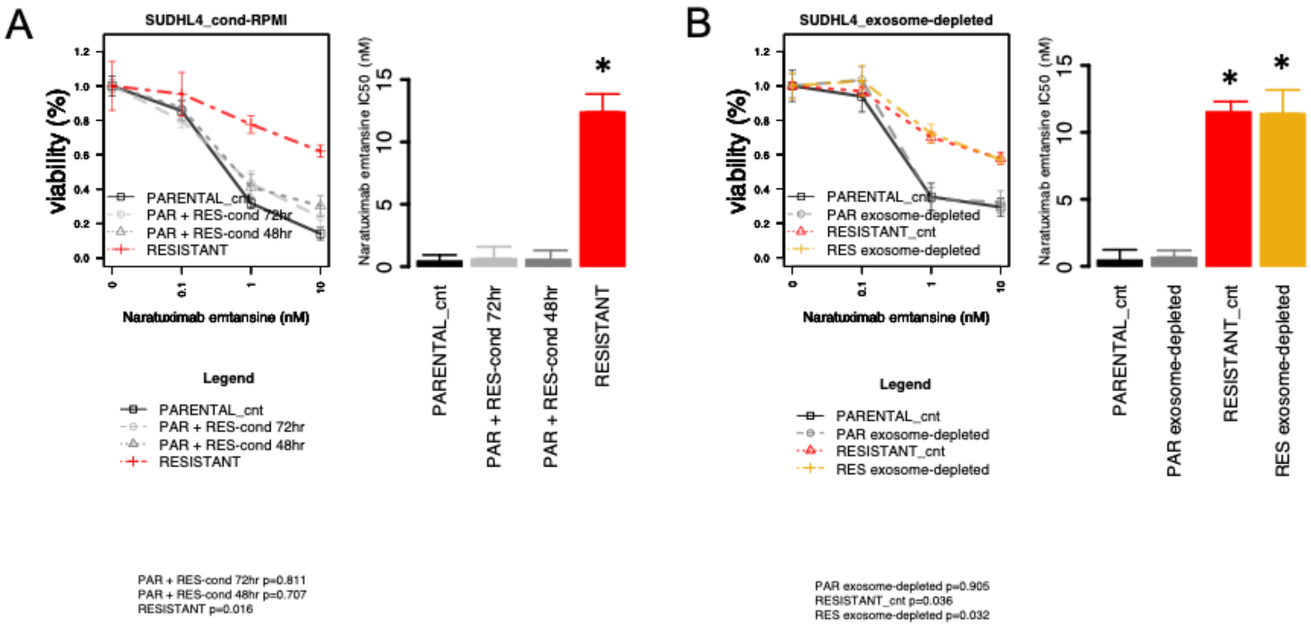

**Supplementary Figure S8. Changes in CD20 surface expression and sensitivity to naratuximab emtansine.** CD20 expression by flow cytometry on SU-DHL-2 (A) and SU-DHL-4 (B) parental and naratuximab emtansine resistant cells. SU-DHL-4 cells with resistance to naratuximab emtansine are also resistant to the combination of naratuximab emtansine plus rituximab (C) and to rituximab as single agent (D). Heatmaps in (left panel, C) represent cell viability upon naratuximab emtansine and rituximab combination, boxplot (right panel, C) shows Chou-Talalay combination index. Barplot in left panel of (D) represents IC50 values of rituximab as single agent. *, P< 0.05. (E) Representative PI staining by FACS obtained in naratuximab emtansine resistant and parental SU-DHL-4 exposed to PBS (control, grey), naratuximab emtansine (200pM, blue), rituximab (10nM, red), combination of the two (green).

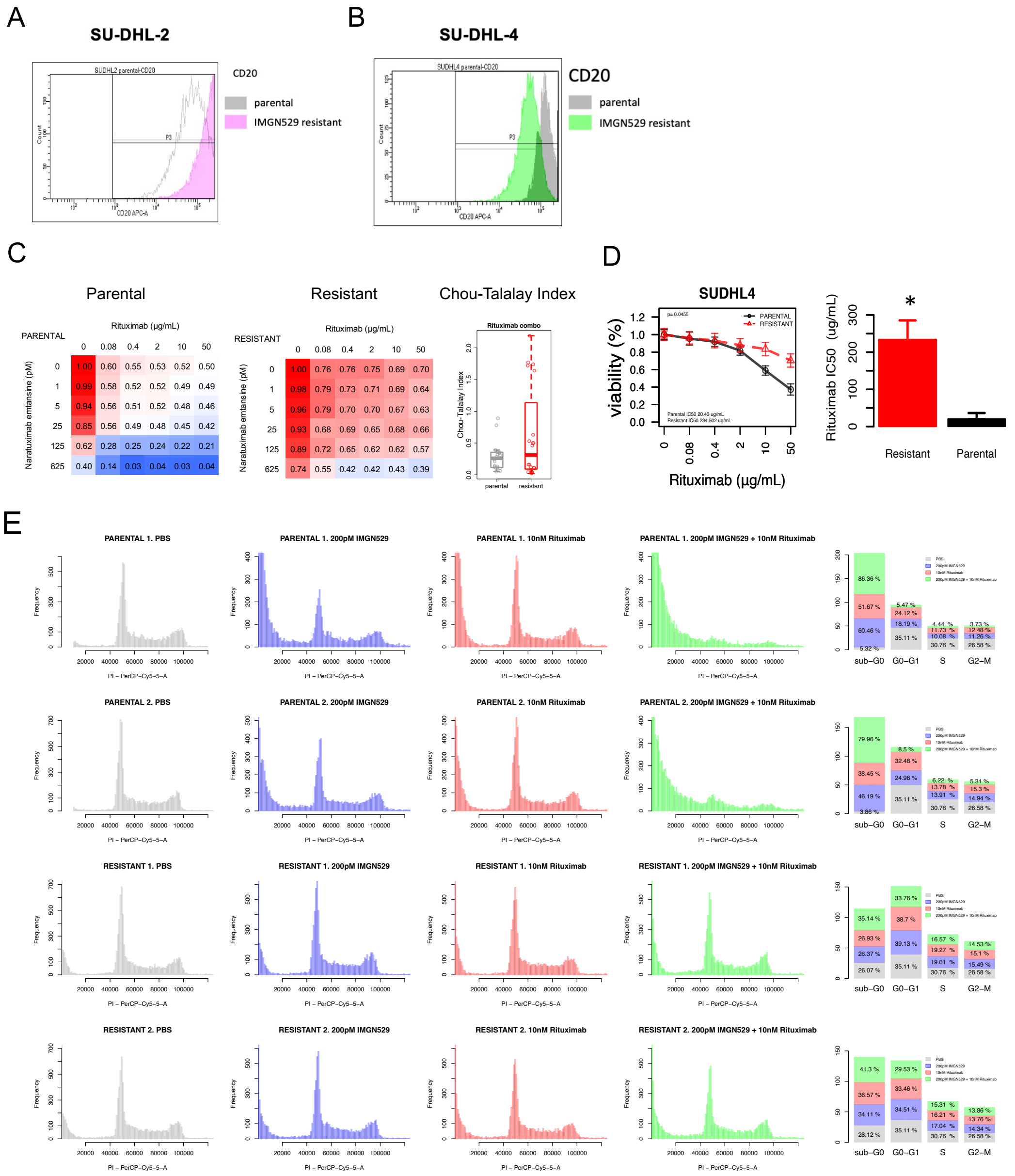

**Supplementary Figure S9. Loss of CD37 expression in naratuximab emtansine-resistant SU-DHL-2 due to a homozygous deletion at *CD37* gene locus in 19q13.33.** (A) Homozygous deletion estimated from WES. Detailed view on 19q13.33. Most frequent splicing variants in blue; green, blue and red bars for single nucleotide variants). (B) Levels of CD37 were evaluated by real-time PCR in parental (grey) and resistant lines of SU-DHL-2 (blue) and SU-DHL-4 (red). Mean of two independent experiments, error bars represent standard deviation of the mean. *, P< 0.05.

A

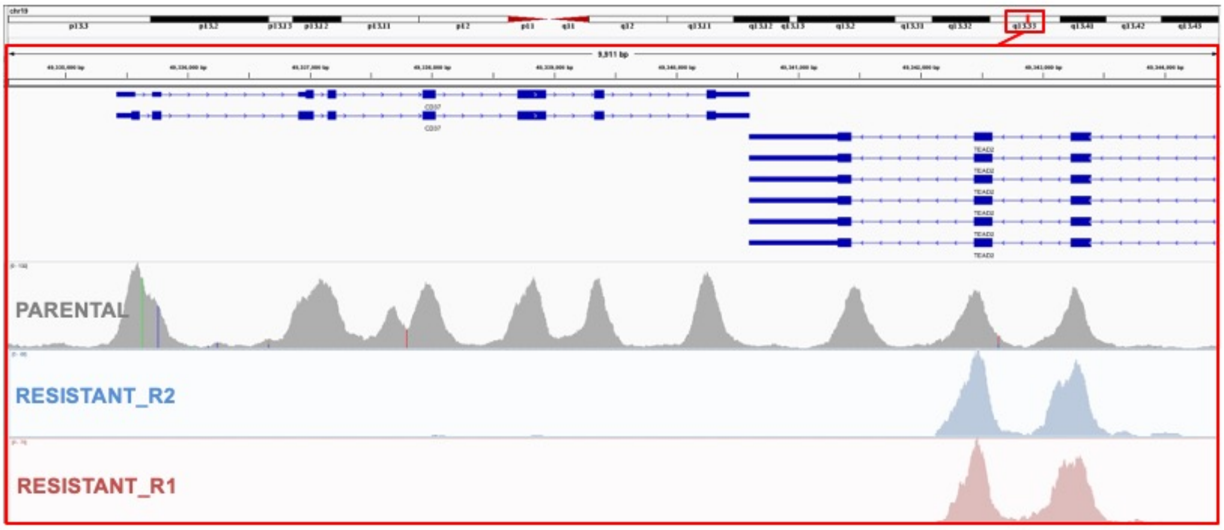


B

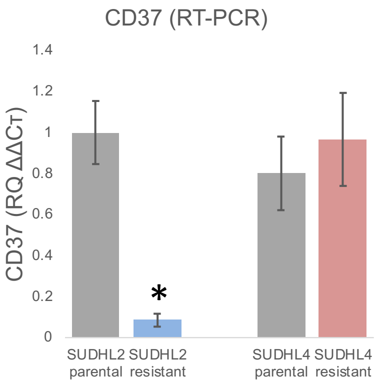

**Supplementary Figure S10. SUDHL4 resistant enrichment pointed to PI3K signaling, lipid metabolism and cell death.** Enrichment map was generated from the output of GSEA (gene set enrichment analyses) comparing parental (blue) and resistant (red) gene expression profiles of SU-DHL-4 lines, using the EnrichmentMap plug-in in Cytoscape (https://apps.cytoscape.org/apps/enrichmentmap).

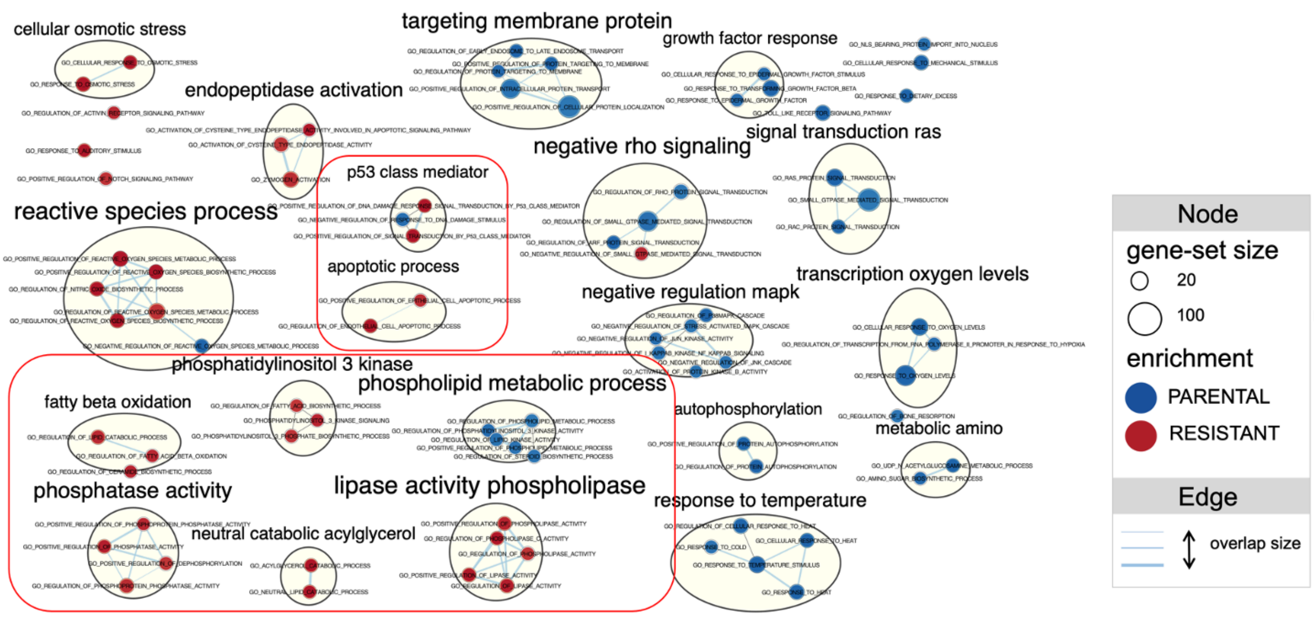

**Supplementary Figure S11. SU-DHL-4 resistant cells exhibited de-regulation of BCL2-family genes.** Gene expression profiles of BCL2 family genes from two independent replicates of resistant compared to parental lines of SU-DHL-4. X and Y axis represents the fold change between each resistant replicate vs parental. Functional annotation represented in the color scheme: anti-apoptocic (blue), oncogene (red), pro-apoptotic (green) and tumor suppressor (orange). Cross for p<0.05 and circle for not significant differences between parental and resistant from a moderated t-test.

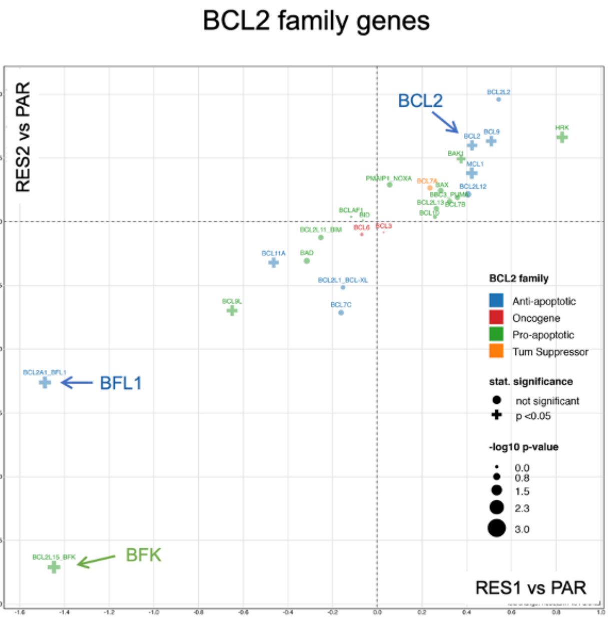

**Supplementary Figure S12.** Structural model of the PI3Kδ complex of p110δ/p85α with the locations of the *PIK3CD* N334T mutation (A) and of other *PIK3CD* mutations recurrent in individuals affected by the APDS as well in DLBCL clinical specimens (B-F).

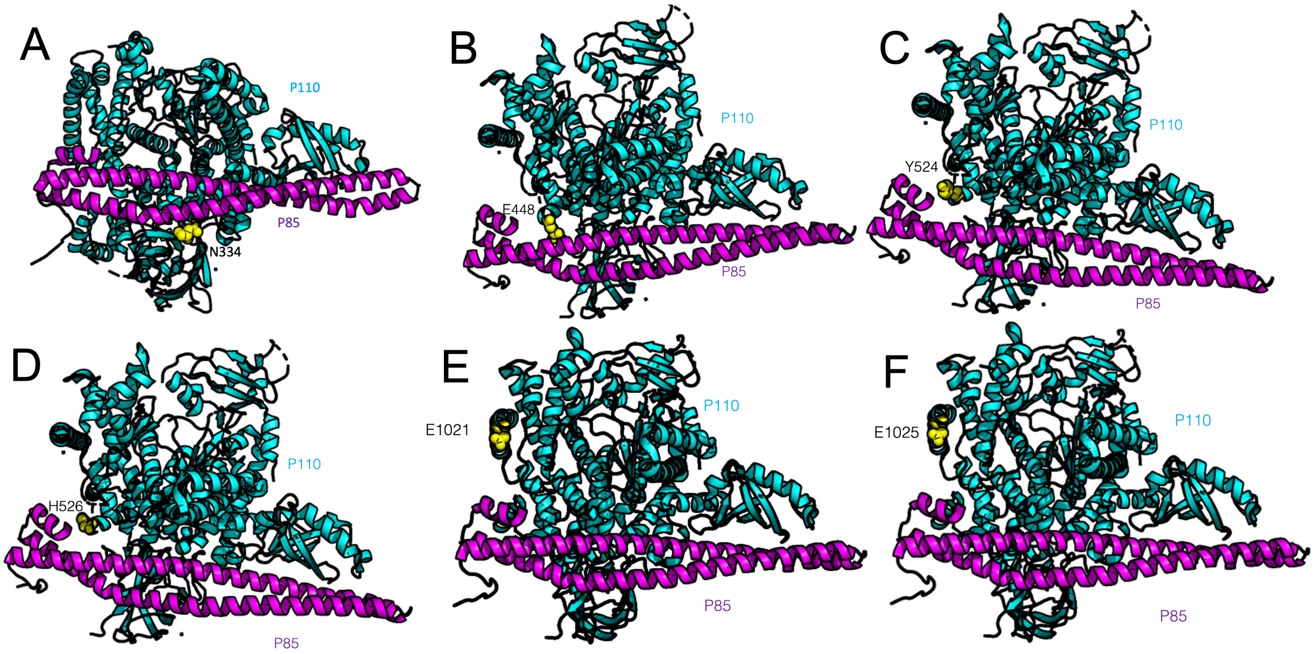

**Supplementary Figure 13. Pretreatment with idelalisib recivers sensitivity to naratuximab emtansine resistant in the resistant SU-DHL-4 cells.** (A) Upper panels show MTT results obtained in naratuximab emtansine resistant and parental SU-DHL-4 (two replicates each) exposed for 72 hours to increasing concentrations of naratuximab emtansine after 72 hours pretreatment with DMSO (left panel), with 100 nM (middle panel) or 500 nM (right panel) of idelalisib. Barplots represents the $IC_{50}$ values. *, P<0.05. (B) Lower-left panel summarizes $IC_{50}$ values obtained in the experiments shown in the upper panel. (C) Representative PI staining by FACS obtained in naratuximab emtansine resistant and parental SU-DHL-4 (two replicates each) exposed to DMSO (control, grey), naratuximab emtansine (625pM, blue) or idelalisib (625nM, red).

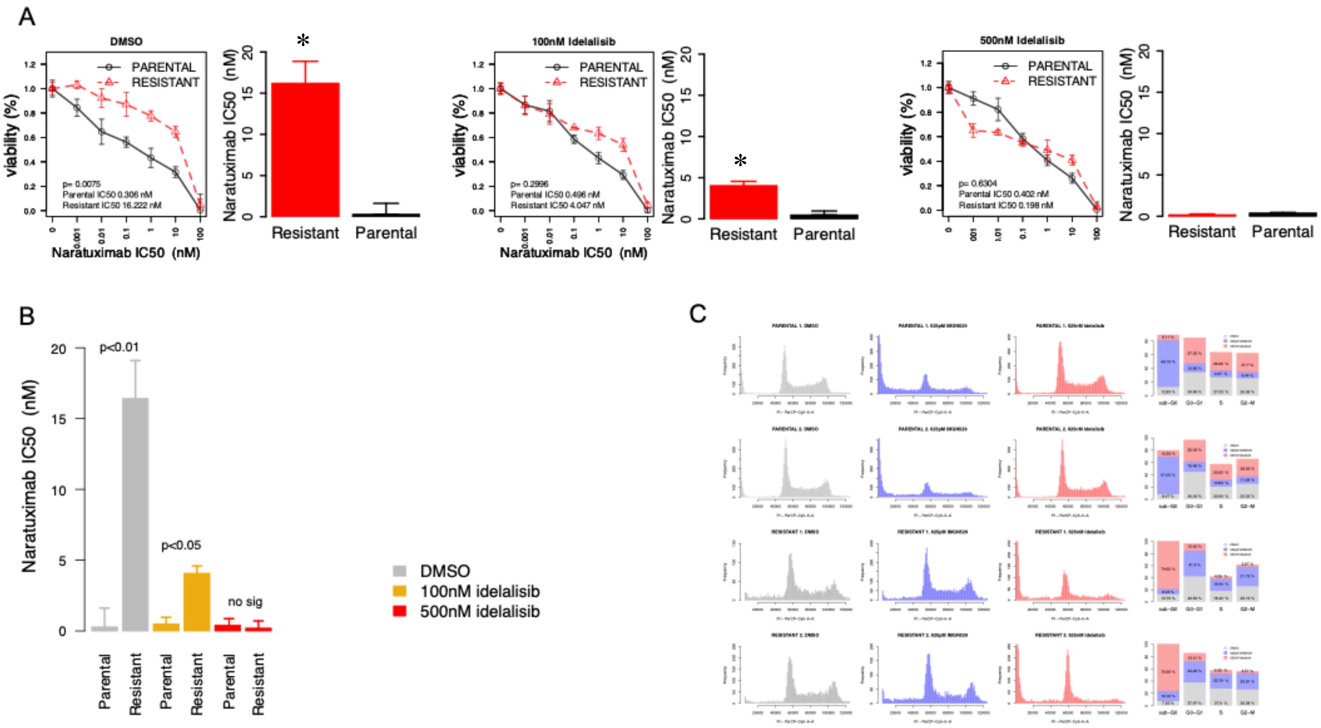


**Supplementary Figure 14.** BH3 profiling of naratuximab emtansine resistant (A) and parental SU-DHL-4 exposed to idelalisib (grey bars) or DMSO (black bars).

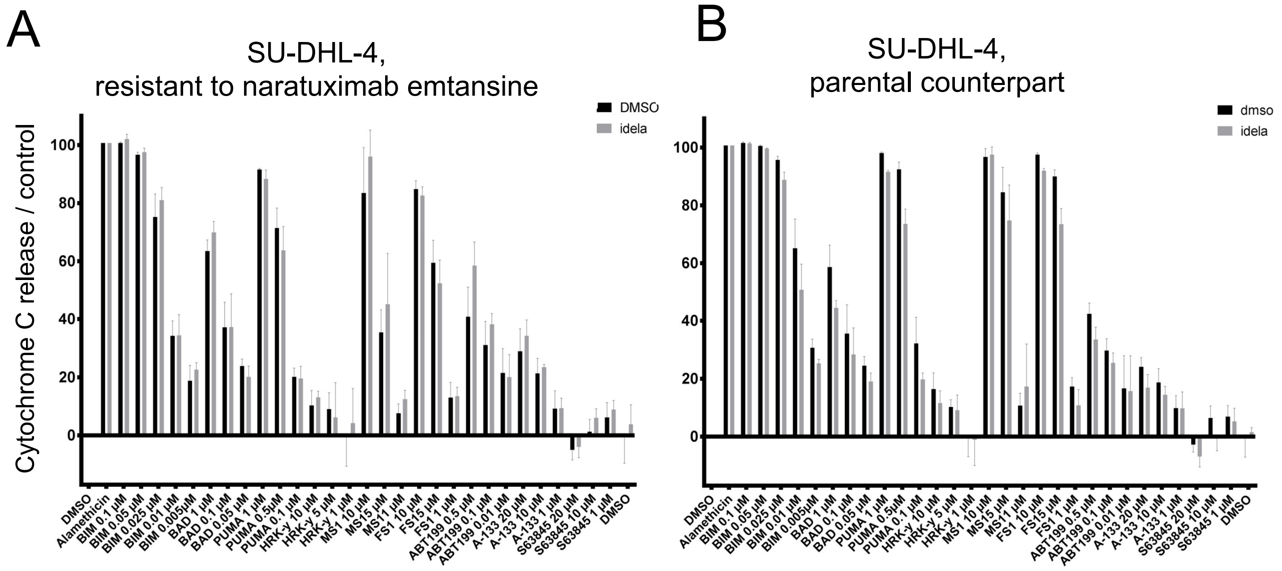
A
SU-DHL-4,
resistant to naratuximab emtansine
DMSO
idela
Cytochrome C release / control
0
20
40
60
80
100
DMSO
Alamethicin
BIM 0.1 µM
BIM 0.05 µM
BIM 0.025 µM
BIM 0.01 µM
BIM 0.005µM
BAD 1 µM
BAD 0.1 µM
BAD 0.05 µM
PUMA 1 µM
PUMA 0.5µM
PUMA 0.1 µM
HRK-y 10 µM
HRK-y 5 µM
HRK-y 1 µM
MS1 10 µM
MS15 µM
MS11 µM
FS1 10 µM
FS15 µM
FS1 1 µM
ABT199 0.5 µM
ABT199 0.1 µM
ABT199 0.01 µM
A-133 20 µM
A-133 10 µM
A-133 1 µM
S63845 20 µM
S63845 10 µM
S63845 1 µM
DMSO
B
SU-DHL-4,
parental counterpart
dmso
idela
0
20
40
60
80
100
DMSO
Alamethicin
BIM 0.1 µM
BIM 0.05 µM
BIM 0.025 µM
BIM 0.01 µM
BIM 0.005µM
BAD 1 µM
BAD 0.1 µM
BAD 0.05 µM
PUMA 1 µM
PUMA 0.5µM
PUMA 0.1 µM
HRK-y 10 µM
HRK-y 5 µM
HRK-y 1 µM
MS1 10 µM
MS15 µM
MS11 µM
FS1 10 µM
FS15 µM
FS1 1 µM
ABT199 0.5 µM
ABT199 0.1 µM
ABT199 0.01 µM
A-133 20 µM
A-133 10 µM
A-133 1 µM
S63845 20 µM
S63845 10 µM
S63845 1 µM
DMSO

**Supplementary Figure S15. Enrichment of the SU-DHL-4 naratuximab emtansine resistant signature in the transcripts higher in DLBCL models with resistance to naratuximab emtansine than in those sensitive to the ADC.** GSEA plots for the enrichment of the SU-DHL-4 naratuximab emtansine resistant signature in baseline gene-expression profiles of DLBCL cell lines that were highly sensitive to naratuximab emtansine (IC50 < 800 pM) compared to the resistant cell lines (IC50 > 10 nM). Green line, enrichment score; bars in the middle portion of the plots show where the members of the gene set appear in the ranked list of genes; Positive or negative ranking metric indicates, respectively, correlation or inverse correlation with the profile; NES, normalized enrichment score.

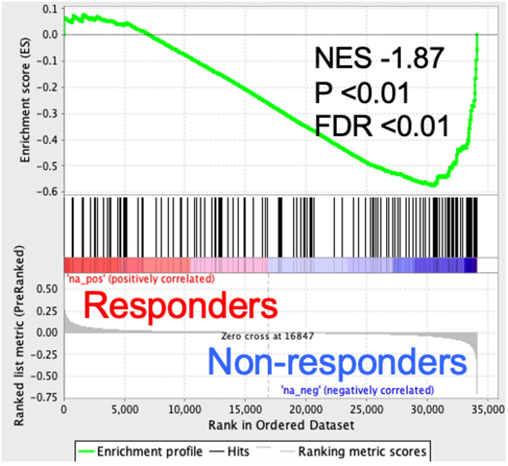

**Supplementary Figure S16. Enrichment of signatures of sensitivity to PI3K inhibitors in the transcripts higher in the SU-DHL-4 naratuximab emtansine resistant than in the parental cells.** GSEA plots for the enrichment of signatures of sensitivity to PI3K inhibitors in the SU-DHL-4 naratuximab emtansine resistant compared to the parental cells. Green line, enrichment score; bars in the middle portion of the plots show where the members of the gene set appear in the ranked list of genes; Positive or negative ranking metric indicates, respectively, correlation or inverse correlation with the profile; NES, normalized enrichment score.

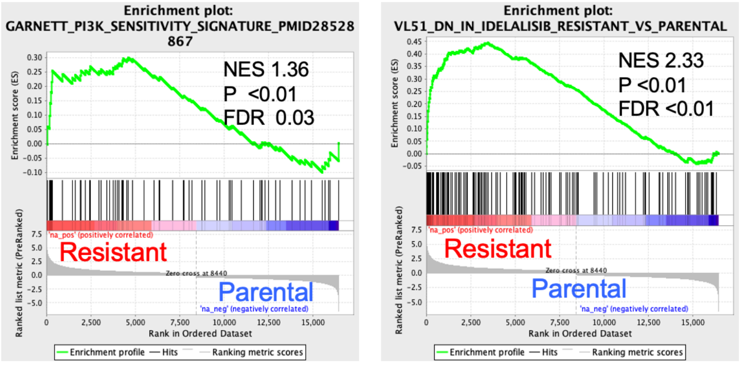


GSEA on SUDHL4 clones comparing resistant vs parental to IMGN529/debio1562

**Supplementary Figure S17. Sensitivity to the PI3Kδ inhibitor idelalisib and to naratuximab emtansine was not correlated.** Pearson correlation between $IC_{50}$ values obtained exposing B cell lymphoma cell lines to naratuximab emtansine and to idelalisib.

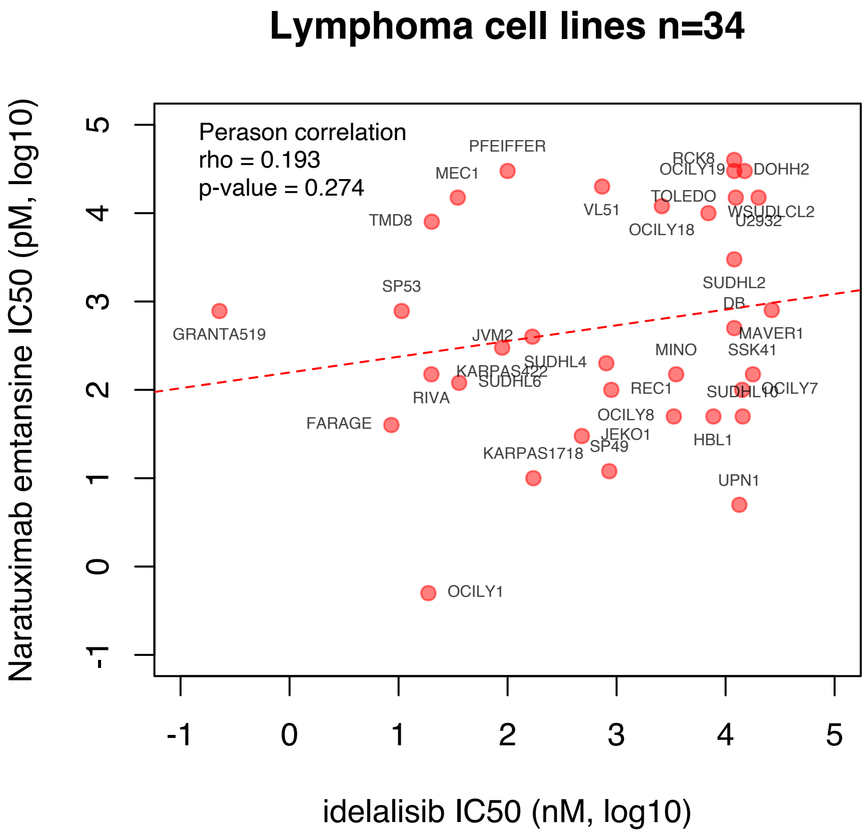
